# Discovery and Development of Novel DNA-PK Inhibitors by Targeting the unique Ku-DNA Interaction

**DOI:** 10.1101/2020.08.24.261875

**Authors:** Navnath S. Gavande, Pamela S. VanderVere-Carozza, Katherine S. Pawelczak, Tyler L. Vernon, Leslyn A. Hanakahi, Matthew Summerlin, Joseph R. Dynlacht, Annabelle H. Farmer, Catherine R. Sears, Nawar Al Nasrallah, Joy Garrett, John J. Turchi

## Abstract

DNA-dependent protein kinase (DNA-PK) plays a critical role in the non-homologous end joining (NHEJ) repair pathway and the DNA damage response (DDR). DNA-PK has therefore been pursued for the development of anti-cancer therapeutics in combination with ionizing radiation (IR). We report the discovery of a new class of DNA-PK inhibitors that act via a novel mechanism of action, inhibition of the Ku-DNA interaction. We have developed a series of highly potent and specific Ku-DNA binding inhibitors (Ku-DBi’s) that block the Ku-DNA interaction and inhibit DNA-PK kinase activity. Ku-DBi’s directly interact with the Ku and inhibit *in vitro* NHEJ, cellular NHEJ, and potentiate the activity of IR and radiomimetics. Analysis of Ku-null cells demonstrates that Ku-DBi’s cellular activity is a direct result of Ku inhibition, as Ku-null cells are insensitive to Ku-DBi’s. The utility of Ku-DBi’s was also demonstrated in a CRISPR gene-editing model where we demonstrate that the efficiency of gene insertion events was increased in cells pre-treated with Ku-DBi’s, consistent with inhibition of NHEJ and activation of homologous recombination to facilitate gene insertion. These data demonstrate the discovery and application of new series of compounds that modulate DNA repair pathways via a unique mechanism of action.

## INTRODUCTION

DNA-PK, a serine/threonine protein kinase, is a member of the PI-3 kinase-related-kinase (PIKK) superfamily that includes ATM, ATR, and mTOR (1). The DNA-PK holoenzyme consists of the 469 kDa catalytic subunit, DNA-PKcs, and the Ku 70/80 DNA binding complex. DNA-PKcs and Ku 70/80 are involved in multiple pathways of DNA metabolism that could impact the development and treatment response of tumors, NHEJ, DDR and telomere stability (2). DNA-PK is required for NHEJ-catalysed repair of DSBs (3). Following the induction of a DSB, the Ku protein binds to the DNA termini; this is followed by DNA-PKcs binding. Auto-phosphorylation of DNA-PKcs and phosphorylation of other target proteins requires assembly of the DNA-PKcs-Ku complex at the DNA terminus which promotes processing, DNA-PK dissociation and the eventual association of the DNA-ligase IV/XRCC4/XLF complex to catalyse the final end-joining repair reaction. DNA-PK also participates in the larger DNA damage response (DDR). In this capacity it can phosphorylate ATM and other downstream targets as well as be phosphorylated by ATM and ATR. This collective response ensures cells can respond to all types of DNA damage to promote survival. Inhibition of the DDR is being pursued with small molecule inhibitors developed to target nearly every kinase in the pathway. Finally, the Ku protein has been localized to telomeres where it is protects the chromosome termini. While the exact mechanisms of protection are still being elucidated, current models include protection from nuclease-catalysed end degradation and more recently, blocking telomere excision and/or recombination(4;5). Importantly, the impact on telomeres is Ku-specific and is independent of DNA-PKcs. Telomere maintenance may therefore be expected to be differentially impacts by Ku inhibitors versus DNA-PKcs targeted agents.

DNA-PK is dysregulated in many cancers, and tumors become reliant on increased DNA-PK activity to respond to the genomic instability associated with unregulated cancer cell growth (6–8). The enhanced ability of tumor cells to repair DSBs is also a major contributor to chemo- and radiotherapy resistance. Thus, blocking DNA-PK dependent repair has been demonstrated in numerous systems to increase sensitivity to both cancer treatment modalities (9–13). Most DNA-PK inhibitors target the active site of the catalytic subunit DNA-PKcs and initial molecules lacked specificity, were metabolic unstable and possessed poor pharmacokinetic profiles that hampered initial development of these molecules (14; 15). More recently many of these liabilities have been addressed and clinical studies have advanced with Merck’s (M3418/nedisertib) and Celgene’s (CC-115) DNA-PK inhibitors being investigated as monotherapy and in combination with IR in the treatment of advanced solid tumours.

Herein we describe the discovery of highly potent and selective DNA-PK inhibitors whose mechanism of action is via blocking the Ku70/80 heterodimer interaction with DNA. We demonstrate a direct interaction of our compounds with the Ku heterodimer and inhibition of DNA binding and DNA-PKcs catalytic activity. We show potent inhibition of NHEJ catalysed ligation activity *in vitro,* in NHEJ-competent whole cell extracts, as well as decreased cellular NHEJ. Data is presented demonstrating the cellular response is a result of Ku inhibition and that the compounds potentiate cellular sensitivity to DSB-inducing agents. In addition, chemically-targeted Ku inhibition improves the frequency of homology-directed repair (HDR) in CRISPR-Cas9 gene editing technology, thus providing new insights and utility of Ku-DBI’s.

## MATERIAL AND METHODS

### Chemistry

The synthesis schemes, procedures and characterization of the compounds are provided in the Supplementary Methods

### Computational methods

Docking studies were conducted using existing crystal structures of the Ku70/80 heterodimer (PDB: 1JEQ) and the Ku70/80 dimer bound to a 55-nucleotide DNA substrate (PDB:1JEY) obtained from the Protein Data Bank (PDB) and prepared them using the Protein Preparation Wizard (16). Force field atom types and bond orders were assigned, missing atoms were added, tautomer/ionization states were assigned, water orientations were sampled, Asn, Gln, and His residues were flipped to optimize the hydrogen bond network, and a constrained energy minimization was performed. Ku-DBi’s were drawn in ChemDraw as MDL molfiles and prepared for docking using LigPrep including a minimization with the OPLS3 force field. All chiral centers were retained as specified in the literature. One low energy ring conformation per compound was generated. Ionization states and tautomer forms were enumerated at pH 7.0 ± 2.0 with Epik.

Ku-DBi’s were flexibly docked into the cleft defined by residues using the Glide Extra-Precision (XP) protocol with default settings (17). Docking poses were evaluated based on visual interrogation and calculated docking score. The poses were selected according to the binding mode and the XP GScore. The Glide Extra-Precision (XP) scoring function was used for the potential amino acid interactions were determined based on proximity to each compound as revealed by docking analysis. The Ku 70/80 interactions with Ku-DBi’s were viewed using Pymol with cartoon, surface, and compounds interaction views. All the molecular modelling within this study was performed using Maestro software, version 11 (Schrödinger) operating in a Linux environment.

### Protein Purification

The Ku 70/80 heterodimer was purified from baculovirus-infected SF9 cells as previously described (18). DNA-PKcs was purified from HeLa cells or HEK 293 cells as described previously (19).

### Biochemical activity assays

Electrophoretic mobility shift assays (EMSAs) were used to assess Ku-DNA binding activity and the effect of potential inhibitors. Reactions were performed with a [^32^P]-labelled 30-bp duplex DNA and purified Ku. The DNA substrate was prepared by annealing 5’-[^32^P]-CCCTATCCTTTCCGCGTCCTTACTTCCCC-3’ to its complement. Ku was pre-incubated with varying concentrations of compound before addition to the EMSA reaction mixture containing DNA. Reactions were incubated for 30 minutes at 25°C and products separated by electrophoresis on a 6% non-denaturing polyacrylamide gel. Gels were imaged using a PhosphorImager and quantified by ImageQuant (Molecular Dynamics) as we have described previously (5;19). Kinase assays to assess DNA-PK catalytic activity were performed measuring the DNA-dependent transfer of [^32^P] from ATP to a synthetic p53-based peptide substrate as described previously (19;20). Briefly, compounds were pre-incubated with DNA-PKcs/Ku for 15 minutes and reactions initiated by the addition of ATP, DNA and peptide. The same 30 bp duplex DNA was used in kinase assays except without the [^32^P] label. The DNA Kinase assays were performed at 37°C for 15 minutes and stopped with 30% acetic acid. Reaction products were spotted on P81-filter paper that was then washed 5 times for 5 minutes each in 15% acetic acid, once in 100% methanol and allowed to dry. Filters were exposed to PhosphorImager screen and analysed using ImageQuant software (Molecular Dynamics). Inhibitor titration assays (both EMSA and DNA-PK) were performed in triplicate and data fit to standard binding curves using Sigmaplot® to calculate IC_50_ values. Autophosphorylation reactions were performed essentially as we have described (21). Briefly, DNA-PK was incubated as described above except omitting the peptide and radioactive tracer. Reactions were terminated by the addition SDS sample buffer and heated to 70oC for 5 minutes. Samples were separated by SDS PAGE on 4-20 % acrylamide denaturing gels and transferred to PVDF. Blots were blocked and probed with the DNA-PKcs phospho-Ser 2056 cluster specific Rabbit monoclonal antibody, and secondary HRP conjugated goat-anti rabbit antibody. Images were captured using chemiluminescence detection. Blots were stripped, re-probed for total DNA-PKcs using a mouse monoclonal and goat anti mouse-HRP secondary and imaged via chemiluminescence.

### DNA Intercalation Fluorescence Displacement Assay (FDA)

The analysis of direct DNA binding of Ku-DBi’s was performed as we have described previously (5). A competitive DNA intercalation assay was performed using SYBR-Green (Sigma) and sonicated salmon sperm DNA (8.29 ng/μL, ThermoFisher). Reactions were carried out in 25 mM MOPS (pH 6.5) and varying concentrations of Ku-DBi’s in black 96-well plates in a final volume of 110 μL. Doxorubicin, a known noncovalent DNA binding chemotherapeutic, was used as a positive control. Fluorescence was measured using a BioTek Synergy H1 hybrid multimode microplate reader with an excitation wavelength of 485 nm, emission wavelength of 528 nm, and a read height of 7 mm. Data were collected using BioTek Gen5 reader software. Reactions were incubated a maximum of 5 min before measurements were collected.

### Thermal-shift assay (TSA)

To directly address the interaction of Ku-DBi’s with the Ku protein we employed a TSA (22). A 220 μL master reaction mixture of Ku with either DMSO as a vehicle control or the indicated Ku-DBi was prepared and 20 μL distributed into 10 individual tubes. The tubes were incubated at the indicated temperatures for 3 minutes in a thermocycler, cooled to 4°C and sedimented at 12,000 x g for 20 minutes. The soluble protein in the supernatant was collected and separated by SDS-PAGE, transferred to PVDF and Ku 70 detected by western blot analysis using chemiluminescent detection. Quantification of band intensities was performed by MultiGage using a Fuji LAS-3000 imager.

### In vitro NHEJ assay

In vitro NHEJ assays were performed essentially as described previously (18). Briefly, reactions (10 μL) were performed in 50 mM HEPES pH 8.0, 100 mM KOAc, 0.5 mM Mg(OAc)2, 1 mM ATP, 1 mM DTT, and 0.1 mg/ml bovine serum albumin (BSA) and the indicated concentration of Ku-DBi. pBluescript DNA was linearized with HindIII and 5’-^32^P-labeled. Reactions contained 10 ng of DNA and 25 μg of whole cell extract. Reactions were incubated at 37°C for 2 h, and terminated by the addition of proteinase K/SDS/EDTA and incubated for an additional 30 minutes at 37°C. Products were separated by electrophoresis on 0.6% agarose gels. Products were detected in dried gels by PhosphorImager (BioRad PMI) analysis.

### Cell culture

Ku80-wt and null mouse embryo fibroblasts (MEF) were originally obtained from Dr. Gloria Li (23) and cultured in RPMI medium supplemented with 10% fetal bovine serum. Cells (5 × 10^3^) were plated in wells of a 96-well plate and incubated for 24 hours prior to any treatments. Cells were then treated with the indicated concentration of Ku-DBi for 48 hours. The vehicle (DMSO) concentration was held constant at 1%. Cell metabolism/viability was assessed by a mitochondrial metabolism CCK-8 assay as we have described previously (24). Following ~~2-hour incubation with CCK-8 reagent, absorbance was measured at 450 nm and compared to vehicle-treated controls to determine percent viability. The results presented are the average and SEM of triplicate determinations.

### NHEJ Host Cell Reactivation Assay

Analysis of cellular NHEJ-dependent repair of DSB lesions was performed using a host-cell reactivation assay as we have previously described (25). Briefly, H460 NSCLC cells in 96-well plates were incubated 24-48 hours prior to analysis. Cells were pre-treated for 2 hours with vehicle or the indicated Ku-DBi in Opti-MEM (Gibco) with DMSO concentration held constant (1%). Cells were then transfected by electroporation using Amaxa nucleofector device (Lonza) with AfeI-linearized pCAG-GFP or unmodified (circular) pMax-GFP (0.2 μg). Complete digestion of the pMax-GFP plasmid (Lonza) was confirmed by agarose gel electrophoresis prior to transfection; the blunt cut site is located between the pMax promoter and GFP expression sites, requiring NHEJ repair for GFP expression. Co-transfection with covalently closed circular pCMV-E2-Crimson (0.1 μg, Takara Bio USA) was used as a transfection control in all experiments. Cells were incubated for 24-48 hours as indicated to allow re-joining of the pMax-GFP. Expression of the fluorescence reporters was identified by flow cytometry (BD FACScan). The percent NHEJ is presented as the ratio of GFP/E2-Crimson compared to control cells transfected with both reporters on covalently closed circular plasmids.

### Cell Irradiation

H460 cells were seeded into 24-well plates 18-24 hours prior to experiments. Two hours prior to irradiation, medium was replaced and fresh medium containing either Ku-DBi or DMSO vehicle was added to cells. Plates were placed on ice for 15 min prior to irradiation. Exponentially growing cells were then irradiated on ice with 2 or 5 Gy of 160 kVp x-rays using a Precision X-ray machine (North Branford, CT) at a dose rate of 67.6 cGy/min. Radiation dosimetry measurements were performed using a Farmer-type ionization chamber (PTW Model N30013, Freiburg, Germany) in conjunction with a Keithley electrometer (Model K602, Cleveland, Ohio). After irradiation, cells were incubated on ice 10 minutes before being allowed to recover at 37° C, 5% CO2 for 24 hours. Viability was assessed by clonogenic survival assays. Cells were detached with trypsin, plated and incubated for 10-14 days. Plates were then fixed, stained and counted. Surviving fractions were determined and normalized to the plating efficiency of cells treated with vehicle only.

## RESULTS

### Optimization of small molecule chemical Ku inhibitors

During our efforts to expand DNA repair targeted therapy, we found the compound **X80** (Figure 1A) displayed modest *in vitro* inhibitory activity when assessing Ku double-stranded DNA binding activity in vitro. We screened a series of ester and carboxylic acid derivatives of the arylpyrazolone moiety (Ring B) and aryl/alkyl ester and amide derivative of Ring A, to identify 5102 and 5135, which displayed dramatically increased potency but both compounds were very insoluble and unstable making the of limited utility for in vitro or cellular analysis (26). Having determined that modifications of X80 could yield more active compounds, we embarked on a multidisciplinary structure-guided synthetic chemistry effort to create high affinity compounds with favorable chemical properties. Utilizing the existing crystal structures of the Ku70/80 heterodimer (PDB: 1JEQ) and the Ku70/80 dimer bound to a 55-nucleotide DNA substrate (PDB:1JEY) we identified a ligand binding pocket for the **X80** core structure located in close proximity to the central DNA-interaction region that surrounds the DNA helix (Figure 1B). The prospective binding pocket spans the interface between the Ku70 (cyan) and Ku80 (magenta) subunits. Closer examination revealed a space-filling pocket located adjacent to ring A of **X80** which spans in the interface between the two Ku subunits (Figure 1C). This interface could accommodate the additional phenyl rings present in the potent 5102 and 5135 molecules (26). We therefore altered the linkage from an ester to an amide to probe this pocket with various **X80** derivatives. The most effective modification we identified was the incorporation of a 3-methoxyphenyl moiety to generate compound **68.** This modification resulted in a 4-fold increase in Ku-DNA binding inhibitory activity (**Figure 2**) and a significant increase in solubility compared to the 5102 and 5135 derivatives (data not shown).

**Figure 1.**
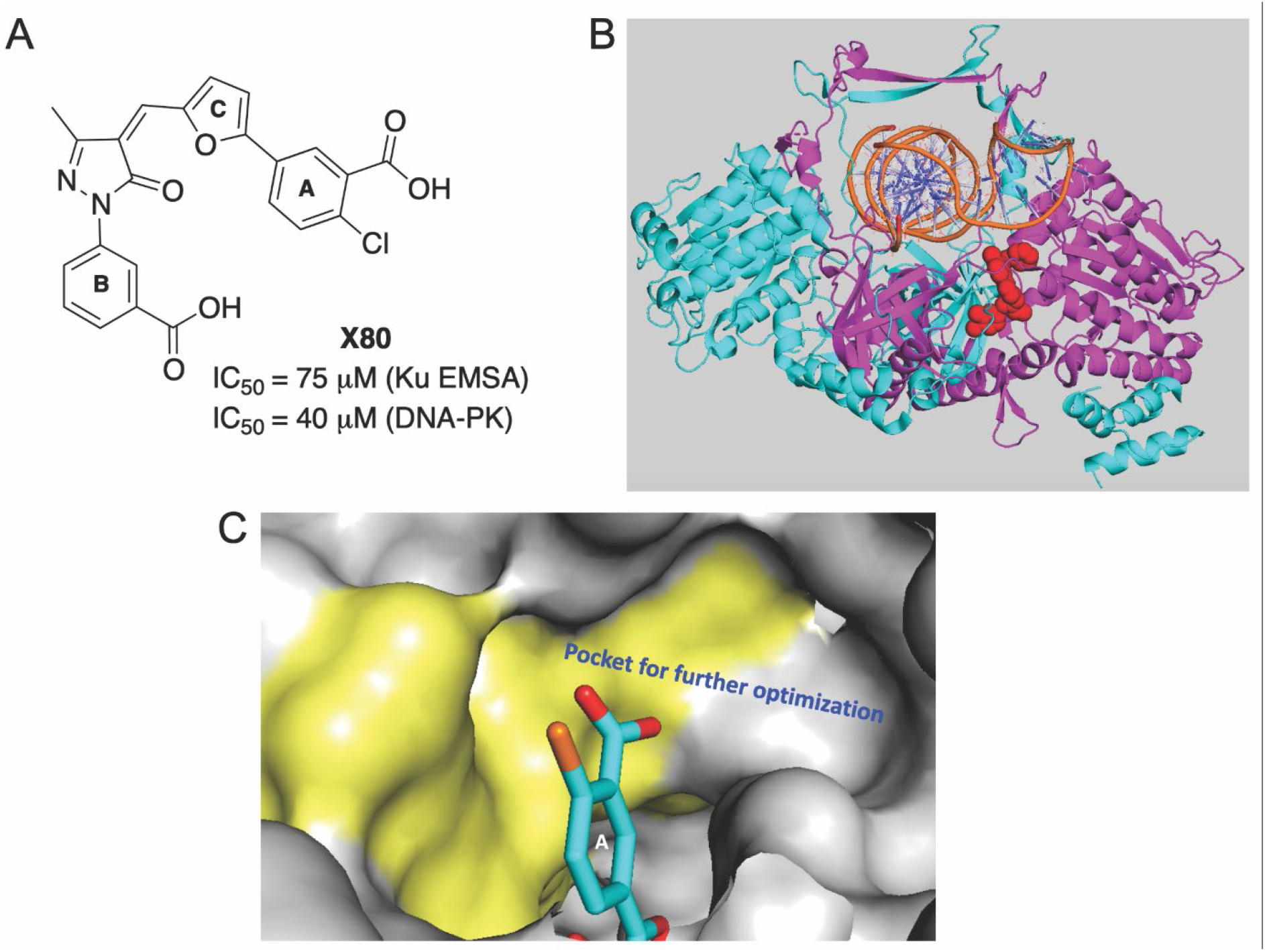
A) Structure of **X80** with Ku EMSA and DNA-PK IC_50_ values; B) Binding pocket of **X80** in the Ku70/80 core. Ku70 is depicted in cyan and Ku80 in magenta. The DNA ring structure is depicted in orange (circular stick model) and **X80** compound in red (spheres); C) Schematic representation of SAR exploration rationale: A pocket surrounding Ring A of **X80** compound for further optimization. (Ku70 shown as yellow and Ku80 as grey surface).

**Figure 2.**
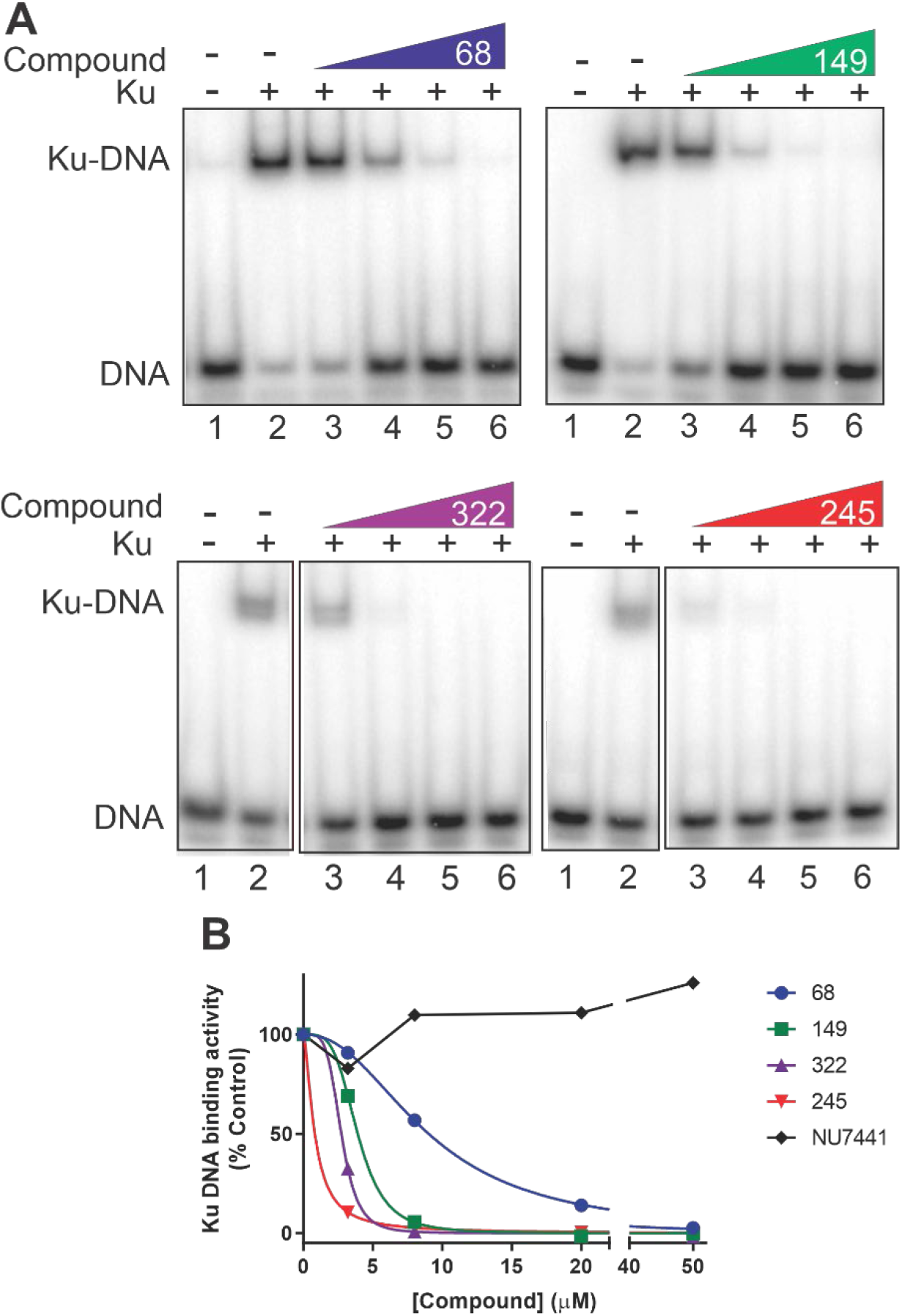
EMSA analysis of Ku inhibitors. A) Inhibition of Ku-DNA binding as assessed by EMSA. Binding assays were performed as described in Methods. The indicated compounds were incubated in reaction with DNA and purified Ku, reaction products were separated by native gel electrophoresis. Gels were visualized and quantified by PhosphoImager analysis. B) Quantification of EMSA data. IC_50_ values were calculated from a minimum of three independent experiments using GraphPad Prism and data presented in Table 1.

The core of the **X80** compound contains carbon-carbon double bond (alkene) between the furan and pyrazolone rings and as such, **X80** derivatives exist as a racemic mixture of *Z and E* isomers with the Z isomer the major product (~73-80%) as determined by NOESY NMR analysis (data not shown). To explore the importance of the double bond we reduced the double bond to a single bond, generating compound **149** which resulted in a further increase in Ku inhibitory activity **(Figure 2**) compared to **68**. The replacement of methyl group of pyrazolone ring in compound **149** with a bioisosteric trifluoromethyl group resulted in compound **322**, and again, a further increase in potency against Ku in a DNA binding assay was observed. These data demonstrate that the double bond is in fact not critical for Ku inhibitory activity and instead is somewhat detrimental to the Ku-inhibitor interaction.

For the final step of optimization, we assessed how bioisosteric modification of the carboxylic acid of phenyl Ring B of compound **149** impacts inhibitory activity. In-silico docking studies showed that carboxylic acid moiety (Ring B) form a hydrogen bond with Arg368 and His243 (Figure 3A) that we posited its importance for Ku inhibition. Interestingly, replacement with a carboxylic acid bioisostere, tetrazole (compound **245**) showed potent Ku inhibitory activity and molecular docking showed more tight binding than carboxylic acid moiety for tetrazole moiety with Ku protein (Figure 3B). The tetrazole moiety and 3-methoxyphenyl group of compound **245** form a hydrogen bond network with a key residue, Arg368. In addition, phenyl-tetrazole moiety is well positioned to make π-π stacking interactions with Phe365. The core structure of compound **149** and **245** were mainly stabilized in a large hydrophobic cavity, hydrogen bond networks, cation-π interaction and π-π interaction formed on the basis of a number of residues spans in the interface between the Ku70 and Ku80 subunits, including Tyr264, Phe365, Ala366, Ala367, Arg368, Asp369, Asp370, Glu371, Ala374, Leu377, Ser 378 (all Ku80 residues) and Tyr416, Glu417, Met446, Pro447, Phe448, Thr449, Glu450, Lys451, Ile452 (all Ku70 residues). Molecular docking (GLIDE) determined similar binding positions for both compounds, with lowest predicted GLIDE XP scores of −7.06 and −7.51 kcal/mol for **149** and **245**, respectively. Both compounds exhibited a good correlation between docking scores and biological activities (Table 1) (17). The 2D interactions of compound **149** and **245** are provided in the Supplementary Data. These data support the importance of the hydrophobic cavity and hydrogen bond network of the carboxylic acid with a tetrazole bioisostere (compound **245**), showed potent Ku inhibitory activity and molecular docking revealed similar tight interactions with the Ku protein (Figure 3B). These data support the importance of the hydrogen bond network in facilitating and/or stabilizing the compound-Ku interaction.

**Figure 3.**
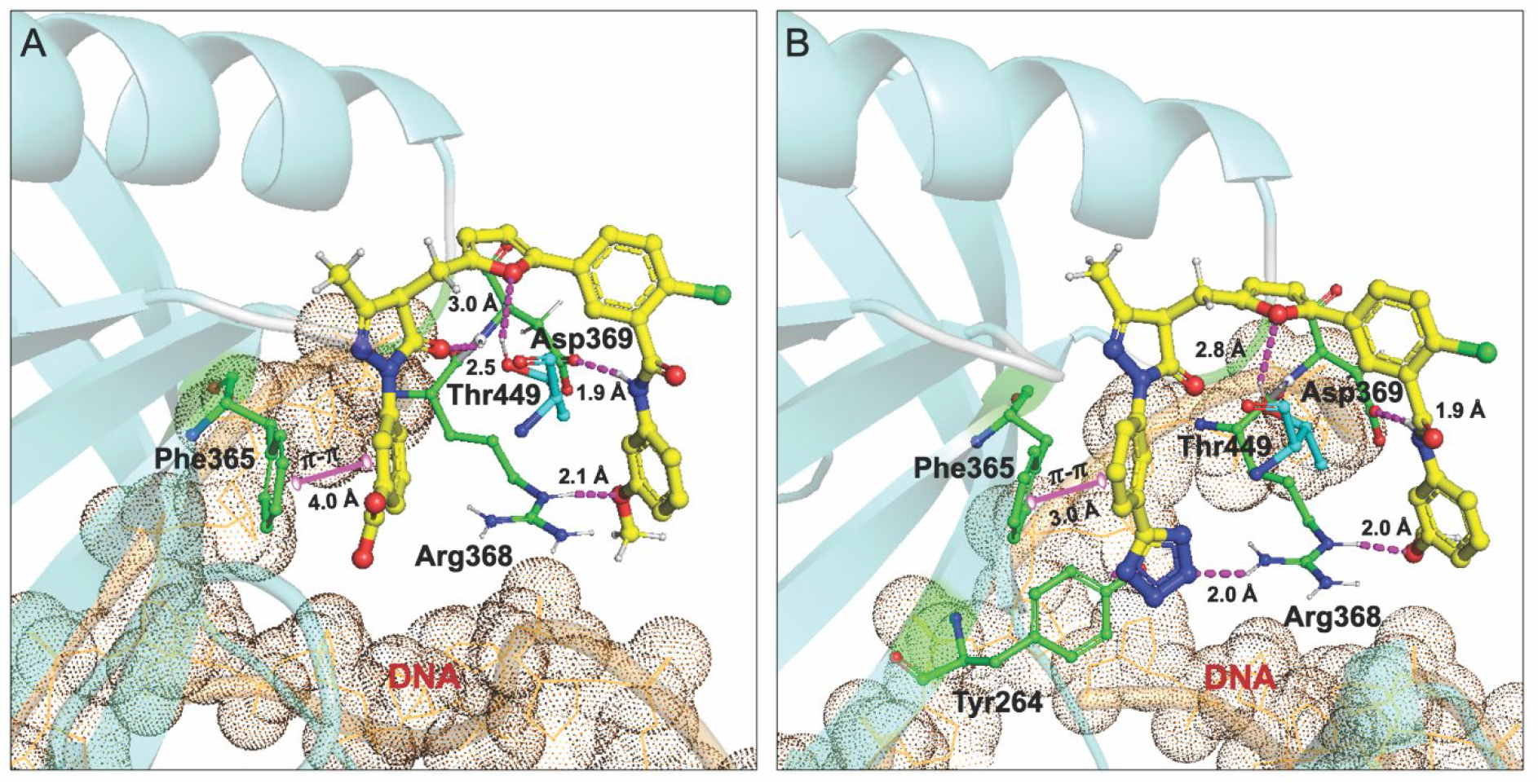
Molecular docking studies (PDB: 1JEQ and 1JEY). A-B) Molecular interactions of compound 149 (A) and 245 (B) (all in yellow carbon) with Ku70/80 heterodimer (key amino acids are shown in cyan carbon (Ku70), green carbon (Ku80) and cartoon is shown in cyan color). The DNA ring structure is depicted in light orange dots. Interaction with amino acid side chains is indicated with the dashed magenta lines and π – π stacking interactions are shown in solid magenta dumbell. Interaction distances indicated in Å.

**Table 1.** Chemical structures and activity profile of Ku-DBi’s

| Assays | 68 | 149 | 322 | 245 |
| --- | --- | --- | --- | --- |
| DNA-PK (IC<sub>50</sub>)<sup>a</sup> | 3.12 ± 0.7 μM | 0.50 ± 0.03 μM | 0.11 ± 0.02 μM | 0.24 ± 0.015 μM |
| Ku70/80 (IC<sub>50</sub>)<sup>b</sup> | 6.02 ± 2.2 μM | 3.72 ± 1.5 μM | 2.66 ± 0.3 μM | 1.99 ± 0.9 μM |
| Glide Gscore<sup>c</sup> | ND | -7.06 | ND | -7.51 |

<sup>a</sup>Determined using DNA-PK phosphoryl-transferase catalytic assay, using a synthetic p53 peptide substrate and highly purified DNA-PK kinase. A commercially available selective DNA-PK inhibitor, **NU7441** showed IC<sub>50</sub> 30 nM in this assay. <sup>b</sup>Compounds were incubated with purified Ku70/80 proteins and direct DNA binding activity measured using EMSA. <sup>c</sup>Predicted molecular docking GLIDE XP Gscore were obtained as described in the **MATERIAL and METHODS** section. ND, Not Determined.

### Mechanism of DNA-PK inhibition

Having established a library of compounds with good stability, solubility and potent Ku inhibition we assessed their effects on DNA-PK kinase activity. *In vitro* kinase assays were performed and the results revealed that the X80 derivatives were potent inhibitors of DNA-PK kinase activity (Figure 4). Interestingly, the IC_50_ values calculated from titration experiments revealed lower values compared to those derived from Ku inhibitory experiments (Table 1). These results are likely a function of the different assays, with Ku binding being a stopped, stoichiometric assay and the kinase assay being a steady-state kinetic assay. The kinase activity data compared favourably to data obtained with the commercially available catalytic site targeted agent, NU7441, which displayed nanomolar IC_50_ values in the kinase assay in vitro while having no impact on the Ku-DNA binding activity (Figure 2B and Figure 4). These results confirm the novel mechanism of action of the Ku inhibitors to inhibit DNA-PK by blocking the Ku-DNA interaction. The in vivo relevant phosphorylation of DNA-PK is the autophosphorylation of the ser 2056 cluster (27)]]. We therefore determine the impact of or Ku-DBi’s on in vitro autophosphorylation of DNA-PKcs via western blot analyses. The data presented in Figure 4C demonstrate that in fact the autophosphorylation of the Ser 2016 cluster is potently inhibited by the individual Ku-DBi’s

**Figure 4.**
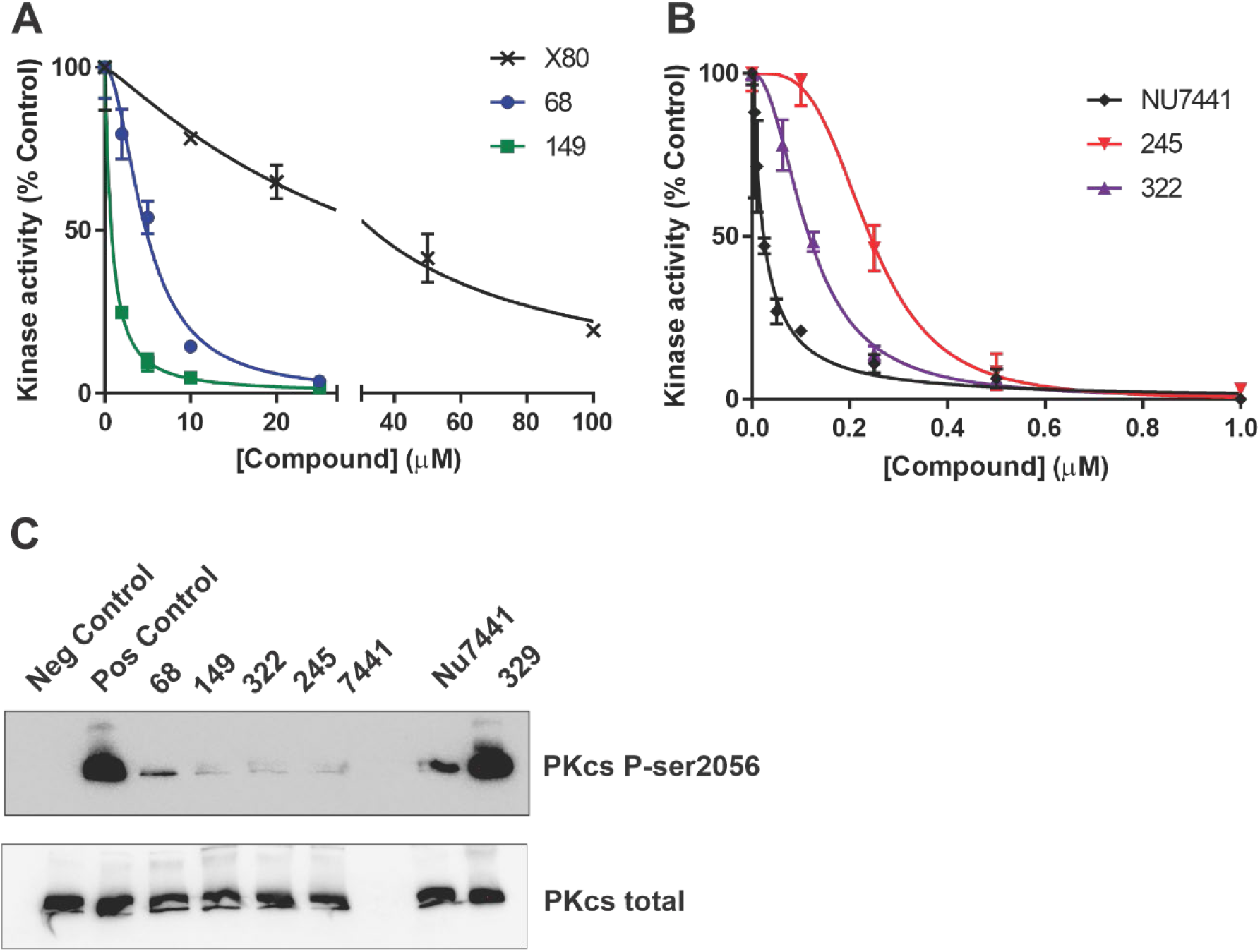
DNA-PK activity is inhibited by Ku targeted agents. A and B). Kinases assays were performed as described in Methods with the indicated compounds. Data are the average and standard error of 2-3 replicates and IC_50_ values were calculated using GraphPad Prism and data presented in Table 1. C) Autophosphorylation of DNA-PKcs at the Ser 2056 cluster was assessed in vitro and detected by western blot analysis as described in Methods. The negative control contained no DNA and ATP and positive control was with DMSO (vehicle). The individual Ku-DBi’s (20 uM) and positive inhibitory control (Nu7441, 5 nM) were included as indicated. Reaction products were separated by SDS PAGE, transferred to PVDF and probed sequentially to detect phospho-DNA-PKcs (upper panel) and total DNA-PKcs (lower panel) as indicated.

While the development of X80 derivatives focused on optimizing direct interactions with the Ku proteins, the mechanism could be a function of the compounds binding to DNA and inhibiting the binding of Ku. We first assessed binding to DNA using a fluorescence displacement assay (FDA) as we have described for the previous X80 derivatives (5;5). Similar to previous results, the data obtained with the newer derivatives show minimal direct DNA interaction up to concentrations 10X the calculated IC_50_’s for kinase activity (data not shown). Therefore, direct DNA binding or intercalation of the Ku-DBi’s cannot account for the observed inhibition of Ku-DNA binding activity and DNA-PK kinase activity. To determine if in fact the X80 derivatives were binding directly to Ku to impart inhibitory activity we employed a thermal stability shift assay (TSA). In this assay the thermal stability of the Ku protein is assessed and the impact of X80 derivative binding on the denaturation profile determined. Briefly, the protein is incubated with vehicle or a fixed concentration of inhibitor and then heated to different temperatures. After heating, the solution is sedimented to precipitate denatured protein. The soluble protein is then separated by SDS-PAGE and detected by Western blot analysis. The data demonstrate that Ku denatures with a Tm of 50°C. The addition of **245** resulted in a 6°C shift in Tm to 56°C (Figure 5A). Together, the FDA and TSA support the conclusion that the X80 derivatives inhibit the Ku-DNA interaction by directly binding to the Ku protein and competitively blocking its association with DNA. The inability to form a functional Ku-DNA complex on the DNA termini leads to reduced DNA-PK catalytic activity by inhibiting the DNA-PKcs-Ku-DNA complex necessary for kinase activity. This represents a novel mechanism of DNA-PK inhibition where kinase activity is abrogated by blocking the Ku-DNA interaction.

**Figure 5.**
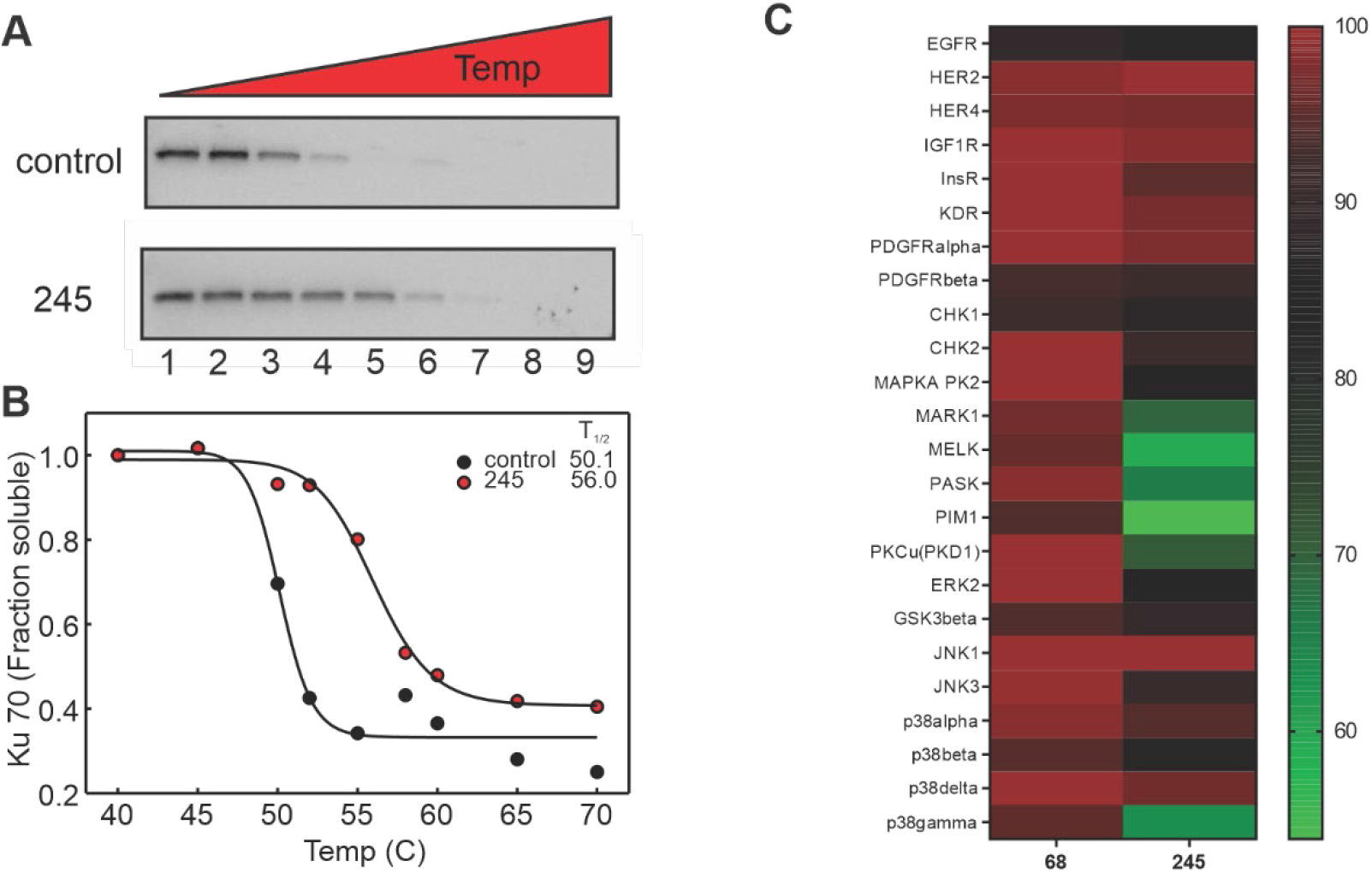
Thermal-shift assay of Ku protein. A) Purified Ku was treated with vehicle or 245 and thermal stability measured by Ku70 western blot. B) Band intensity was normalized to the 40°C point and plotted versus temperature. Data were fit to a 4-parameter sigmoidal curve and T1/2 determined. The error associated with the fits were less than 0.5°C. C) Kinase specificity. The indicated compounds were assessed in a kinase screen using ADP-Glo as described in Methods. The percent activity remaining for each kinase was calculated and presented as a heat map.

### Specificity of Ku-DBi’s

Having established the activity of these compounds in vitro, we sought to assess their specificity. As we originally discovered the X80 class of compounds in a virtual screen based on XPA (Xeroderma Pigmentosum Group A), we determined the cross reactivity of the Ku-targeted X80 derivatives with XPA. No inhibition of XPA-DNA binding activity was observed at concentrations below 20 μM (data not shown). These data also support the novel mechanism of action Ku-DBi’s as a direct DNA binding molecule wold be expected to block other DNA binding proteins. With the explosion of kinase inhibitor development and assays to determine kinase specificity, we assessed a series of kinases for inhibition with Ku-DBi’s. As DNA-PK is the only kinase that requires Ku DNA binding for activity we wold expect minimal inhibitory activity against other kinases. The results demonstrate that the majority of the 24 kinases tested are not appreciably inhibited by the X80 derivatives (Figure 5C). Interestingly, the MELK kinase does show moderate inhibition by the **245** consistent with previous data demonstrating that arylpyrazolone moiety possess considerable affinity for the kinase active site and block kinase activity (28). Similarly, the PIM1 kinase possesses affinity for 3,5-diphenyl substituted indole/oxoindolins that adopt a similar conformation to the **X80** derivatives and amide modification increased the affinity, consistent with the activity of inhibitors **68** and **245** (29;30). Overall these data demonstrate the ability to design potent and specific inhibitors of the Ku-DNA interaction.

### Chemical Ku inhibition blocks NHEJ-catalysed repair of DSBs

To determine how chemical inhibition of Ku impacts NHEJ, we performed an *in vitro* assay to assess the joining of DNA termini catalysed cell-free extracts (25). Briefly, linearized plasmid DNA (3kbp) is incubated with a cell-free extract prepared from an NHEJ-competent cell (HEK 293) and ATP. Data demonstrate that all Ku inhibitors tested (6 out of 6) inhibited *in vitro* NHEJ. The representative experiment shows inhibition of end-joining in a concentration-dependent manner for compound **149** (Figure 6A). These data demonstrate target engagement in a complex protein mixture and inhibition of DNA-PK dependent NHEJ. The relative potency compared to the our EMSA and kinase assay is somewhat reduced which suggests that in the complex cell-free extract the effective concentration of Ku-DBi is reduced possible via low affinity interactions with other proteins or is being degraded. Data supporting the reduced effective concentration in extracts was obtained in analysis of DNA-PKcs autophosphorylation where the potency of Ku-DBi’s was reduce compared to highly purified DNA-PK preparations (data not shown).

**Figure 6.**
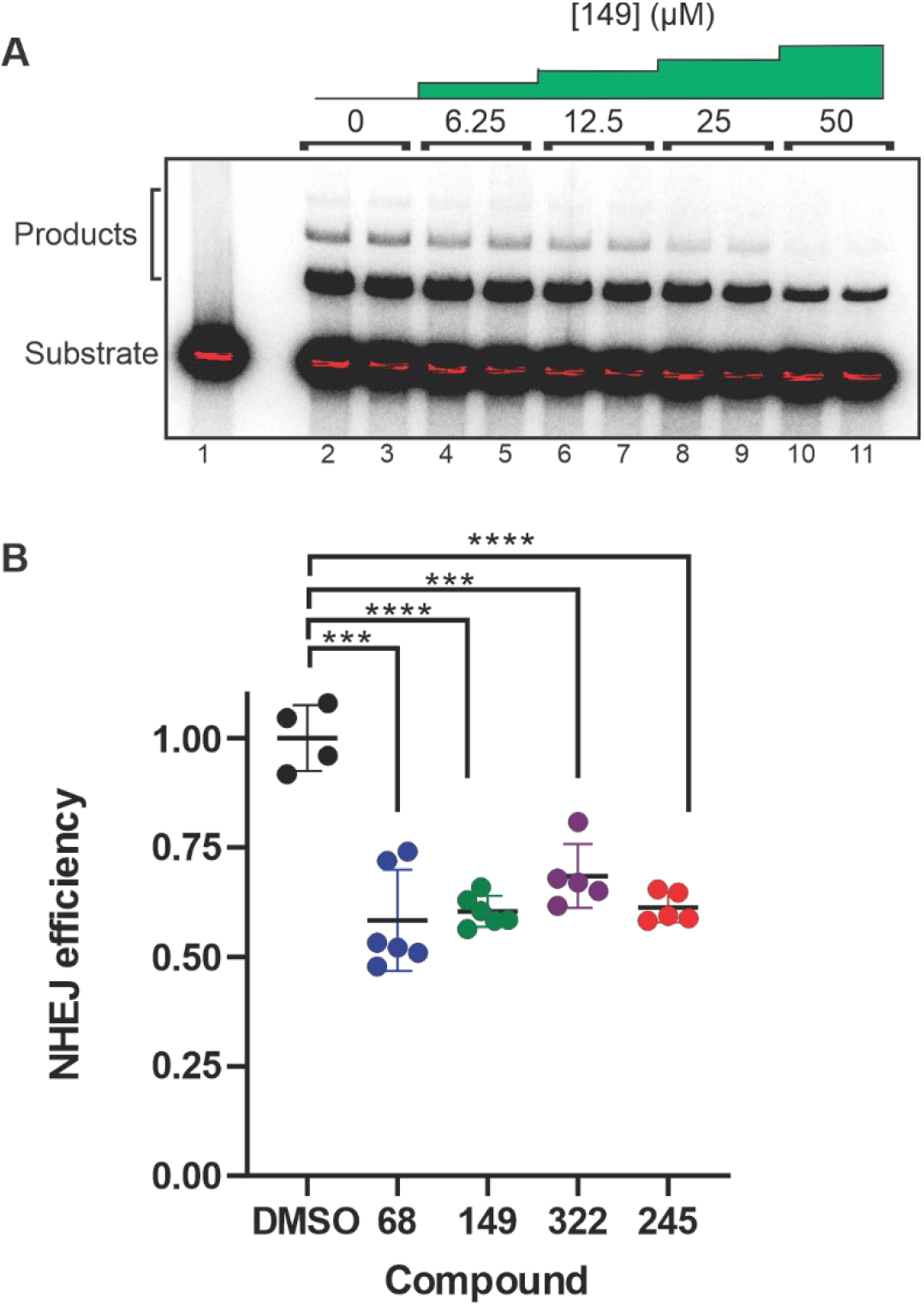
Analysis of Ku targeted DNA-PK inhibitors on *in vitro* and cellular NHEJ. A) A radiolabelled linearized plasmid DNA (3kbp) substrate (Lane 1) and the formation of plasmid multimers when increasing concentration of compound **149** was incubated with the extract (lanes 2-11). Quantification of the data revealed an IC_50_ of ~15 μM for compound **149**. B) Cellular NHEJ was assessed by host cell reactivation as described in Methods. Individual data points are presented along with the average and standard error.

Having demonstrated inhibition of NHEJ *in vitro,* we sought to determine if the chemical Ku inhibitors could abrogate cellular NHEJ. We employed a host cell reactivation assay in H460 NSCLC as we have previously described to measure cellular NHEJ activity (25;31). Briefly, a linearized plasmid encoding a GFP gene was transfected into cells along with a circular RFP expression plasmid as a transfection control. GFP expression is obtained only upon NHEJ-mediated re-circularization of the plasmid. The ratio of green to far-red cells is used for quantification as measured by flow cytometry. We found that incubation in serum-free medium greatly increased the cellular activity of the Ku-DBi’s, again consistent with non-specific protein binding or degradation (not shown). In addition, we observed greater activity in cells pre-incubated with Ku-DBi’s for 2 hours prior to transfection. The data presented in Figure 6B demonstrate that all Ku-DBi’s are capable of significantly reducing NHEJ-catalysed repair events. The data show an approximate 50% reduction in NHEJ catalysed repair events. Interestingly the relative difference between the individual Ku-DBi’s was not evident in this assay suggesting the potential for off-target interaction of the Ku-DBi’s

### Cellular sensitization to DSBs and assessment of on-target activity

Having determined that our Ku inhibitors block cellular NHEJ, we performed the crucial experiment to determine if Ku-DBi’s can sensitize cells to DNA DSB-inducing agents and determine if this effect is the result of a direct on-target interaction between the Ku 70/80 heterodimer and the Ku-DBi. To determine if the chemical inhibition of Ku impacts sensitivity to DNA DSB inducing agents, we assessed cell viability of wild type (wt) and Ku 80-null MEFs to DSB-inducing agents with and without Ku-DBi’s. The data presented in Figure 7A shows cellular viability in cells treated with increasing concentrations of the radiomimetic drug bleomycin with and without Ku-DBi’s. Cells were treated concurrently for 48 hours with 20 μM **68** and the indicated concentrations of bleomycin. We selected 20 μM **68** based on preliminary single agent experiments demonstrating less than a 10% reduction in survival, consistent with limited toxicity in normal cells (not shown). As expected, Ku80-null cells displayed increased sensitivity to bleomycin (~6-fold) compared to the wt MEFs. The increased sensitivity to bleomycin induced by **68** is also readily apparent in the wild type cells (compare black circles to blue circles). Importantly, there is no significant increase in sensitivity induced by **68** in the Ku80-null cells (compare black triangles to blue triangles). These data conclusively demonstrate that the observed sensitization effect of **68** to bleomycin is mediated through on-target inhibition of Ku-DNA binding by the Ku-DBi. Similar results were observed when the effect of **245** on cellular sensitivity to etoposide was assessed (data not shown). The data also demonstrate that chemical inhibition with Ku-DBi **68 or 245** do not reach the sensitivity of the Ku null cells. This indicate there remain the possibility to increase the activity of these inhibitors by either increasing potency or cellular uptake and stability.

**Figure 7.**
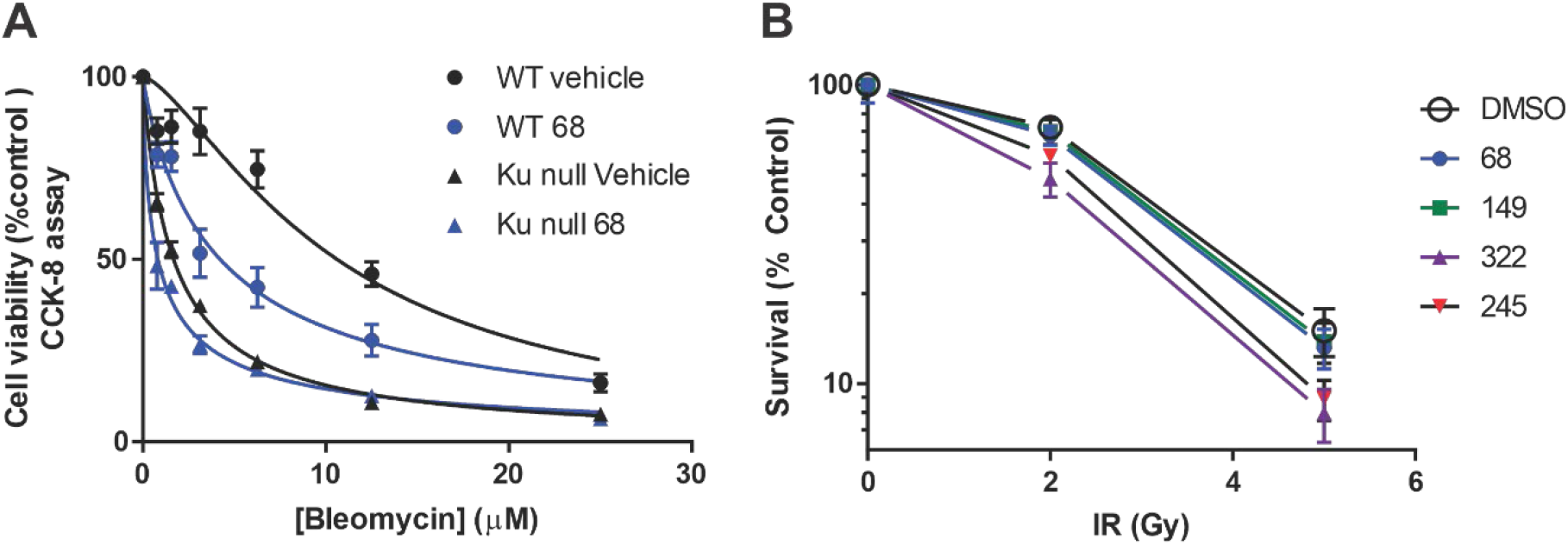
Sensitization to DSB inducing agents by KU targeted DNA-PK inhibitors. A) Cellular inhibition of Ku-DNA interaction by compound **68**. Cells were seeded and treated with vehicle or 20 uM **68** and the indicated bleomycin concentration. Cell viability was assessed by CCK-8 metabolic assay and data are presented as the mean and SEM of triplicate determinations. B) Enhancement of cell killing after irradiation. After irradiation, cells were plated to assess clonogenic survival as described and data represent the mean and SEM of triplicate determinations.

A similar series of experiments were conducted to assess the impact of Ku-DBi’s on radio sensitivity. H460 NSCLC cells were treated in 24 well plates with either vehicle or the indicated Ku-DBi for 2 hours prior to irradiation with either 2 or 5 Gy of X-rays. Cells were allowed to recover for 24 hours then plated for colony formation and determination of clonogenic survival. An enhancement of radiosensitivity was observed in cells treated with the Ku inhibitors. and the relative sensitization follows the in vitro potency for the Ku-DBi’s (Figure 7B).

### Increased efficiency of HDR-mediated gene editing by chemical inhibition of Ku

Recent studies have shown that reduced NHEJ activity *in vivo* results in increases in HDR activity, and this phenomenon can be exploited to increase the efficiency of HDR-mediated CRISPR/Cas9 precision genome engineering (32–35). To investigate the effect of Ku inhibitors on CRISPR/Cas9 genome editing, a donor molecule encoding a GFP and a puromycin resistance gene flanked by 800bp of sequence homologous to a CRISPR cut site in the EMX1 gene was synthesized (Sigma). Donor, gRNA targeted to EMX1 and Cas9 plasmids were co-transfected into H460 cells in the presence and absence of Ku inhibitor **245**. Seventy-two hours after transfection, cells were single-cell diluted in 96-well plates and placed under puromycin selection and cultured until confluent. Cells were harvested and genomic DNA isolated for PCR analysis. Junctional PCR analysis was used to assess targeted insertion of the donor molecule Figure 8. Of the 28 puromycin resistant clones from the vehicle treatment, 4 of them were positive for precise transgene insertion (Table 2). Conversely, 4 out of the 5 puromycin resistant clones derived from the cells treated with 20μM **245** were positive for precise transgene insertion (Table 2). These results indicate that treatment with Ku inhibitor increased the efficiency of HDR mediated gene insertion at a DSB created from CRISPR/Cas9 by 6-fold. The data suggest that the increase in gene insertion efficiency is not necessarily an increase in HR but a decrease in inaccurate integration events. This data suggests that Ku-DBi’s could be effective to reduce off-target, potentially mutagenic events that have hampered Crispr mediated therapeutic applications.

**Figure 8.**
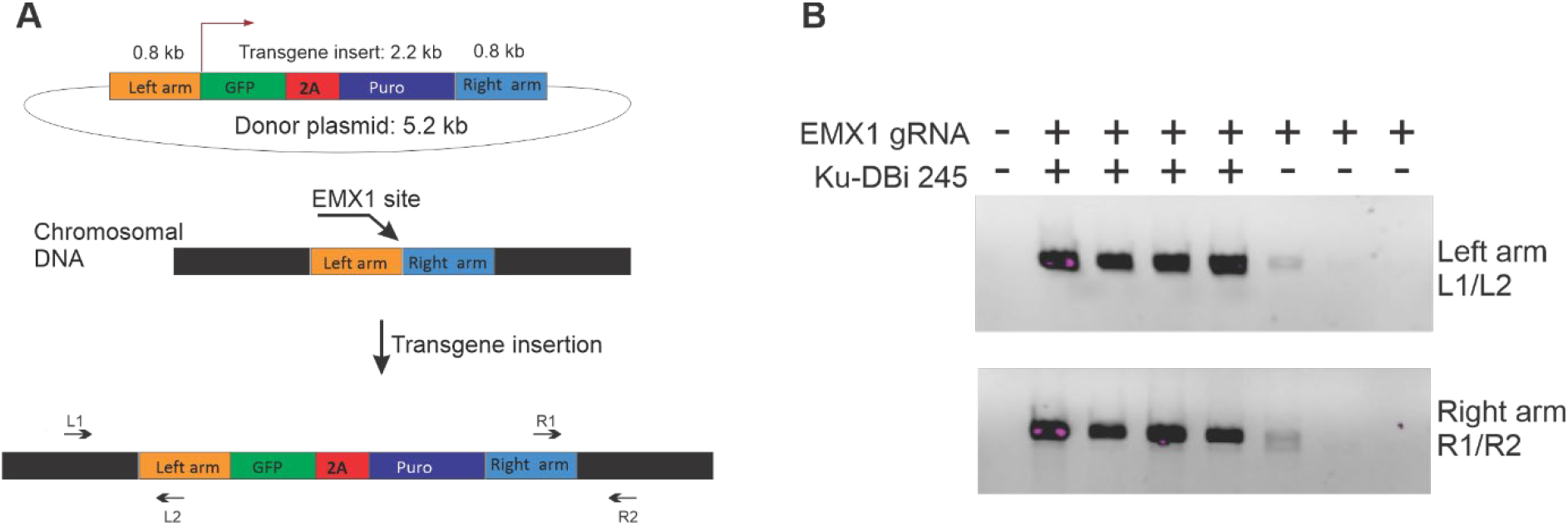
Enhancement of HDR mediated gene insertion. A) Schematic of gene insertion experiment. The 5.2 kbp donor plasmid was designed with a 2.2 kb insert to express GFP and confer puromycin resistance. Homology arms of 800 bp were included on either end to target the insertion to the genomic EMX1 site. PCR primers were designed to allow amplification of the left and right junctions to assess accurate gene insertion B) Genomic DNA was analysed for precise gene insertion by in/out PCR analysis using the primers depicted in Panel A.

**Table 2.** Ku-DBi modulation of CRISPR/cas9 mediated gene insertion

|  | vehicle | Ku-DBi **245** |
| --- | --- | --- |
| Clones isolated | 28 | 5 |
| PCR positive clones | 4 | 4 |
| Insertion efficiency (%) | 14 | 80 |

## Discussion

NHEJ is a predominant repair pathway in human cells, and has historically been considered a guardian of the genome through preventing genomic instability after the occurrence of a DNA DSB. Various genetic studies have revealed that the loss of core NHEJ components, including Ku70/80, DNA-PKcs, DNA Ligase IV, XRCC4, and XLF leads to genomic instability, as evidenced by an increase in chromosomal translocations and aberrations. Defects in key NHEJ components also result in increased sensitivity to DNA damaging agents like ionizing radiation. Despite the importance of this cellular pathway, many questions regarding the intricacies of the molecular mechanisms driving NHEJ remain. One of the biggest challenges in the field remains the determination of the molecular mechanism that drives activation of DNA–PK. Specific roles in NHEJ and V(D)J recombination have been elucidated from both genetic, structural and biochemical analyses that firmly position it as the end recognition complex binding to DNA termini. Abundance and affinity would suggest that DNA-PK would likely be the first protein to arrive at a DSB, a point that has been debated, though recent evidence provides experimental evidence for the rapid association of DNA-PK with DNA ends in living cells (36). Biochemical and structural studies support the hypothesis that DNA–PK activation is driven by direct contact with DNA (37–39) and there is evidence that DNA structure and sequence play a unique and critical role in optimal DNA-PK activation (19;20). Previous work has shown that the Ku heterodimer is required for efficient NHEJ and functions as a scaffold for other NHEJ proteins to bind (40), including forming a critical interaction with DNA-PKcs. Our group has previously shown the Ku80 carboxy-terminus domain (CTD) must be tethered to the Ku core DNA binding domain to support its role in DNA-PK activation (41). The data presented in this manuscript support the importance of Ku-DNA binding in relation to DNA-PK kinase activity, as direct chemical inhibition of Ku-DNA binding activity abolishes kinase activity in a specific fashion (Figure 4 and Figure 5). This conclusion is supported by results utilizing the DNA-PKcs inhibitor (NU7441) that show no inhibition of Ku-DNA binding but similar inhibition of DNA-PK kinase activity (Figure 2B and Figure 3B). Confirming our conclusion that DNA-PK kinase activity is driven by a Ku-DNA interaction that is disrupted by our novel small molecule inhibitors that bind to the Ku protein, we demonstrated a direct inhibitor-Ku interaction (Fig 5A) and failed to detect binding of DBi’s to DNA data not shown). Small molecule inhibitors are anticipated to have great utility to further delineate the molecular mechanisms driving Ku binding and subsequent DNA-PK activation.

The molecular mechanisms that govern DSB repair pathway choice remain enigmatic. Two critical DNA double strand break repair pathways exist (NHEJ and homology directed repair (HDR) and importantly, cell cycle stage plays a key role in the choice of the repair pathway after a DNA double strand break occurs. Despite the general understanding that HDR is active primarily in S and G2 phases of the cell cycle while NHEJ is the predominant mechanism utilized in G1 phase and non-dividing cells, it continues to remain unclear how the cell determines which pathway to use for the repair of a DSB. This is becoming increasingly important, as more recent research has suggested that joining of replication associated breaks can lead to genomic rearrangements and general genomic instability, highlighting the importance of pathway choice for maintaining genome integrity (42). Our novel molecules have the capability to inhibit NHEJ activity in cells (Figure 6) through blocking the first step of the NHEJ pathway, Ku-DNA binding. Having the availability of chemical inhibitors to target specific steps in the pathways will allow the determination of detailed mechanisms of signalling and repair. The advent of DNA-PK inhibitors targeting the kinase activity directly through ATP-mimetics has proven extremely useful to determine global roles in repair, though specificity of earlier molecules obscured specific roles for the DDR kinases. The ability to specifically block these activities can also provide added insight into the biochemical mechanisms of action. The difference in how cells responds to a DSB when DNA-PK is inhibited by a kinase targeted agent is likely to be very different than a molecule that blocks the interaction with DNA. One could envision inhibiting the kinase resulting in a longer residence time of the DNA-PK complex on the DNA termini, as autophosphorylation has been demonstrated to increase the dissociation rate of DNA-PK from the DNA terminus (21;43;44). Longer residence however, could either promote or inhibit recruitment or binding of other proteins at the DNA discontinuity to alter the repair pathway employed. The ability to inhibit the initial DNA-binding event is likely to reveal additional complexities associated with double strand break repair pathway choice as having a protein present but inactive at the termini is surely different than not having anything bound to the terminus.

Exploiting the mechanism of DSB repair regulation to enhance genome engineering remains an active area of investigation. It is well understood in the genome engineering field that increasing the efficiency of HDR would render CRISPR/Cas9 genome engineering a faster, easier and more accurate process. Various studies in the DNA repair field have shown that HDR activity can be enhanced by inhibiting NHEJ (32;45) (46). It has also been shown that inhibiting critical NHEJ proteins through genetic modifications as well as with chemical inhibitors or siRNA can result in an increase in HDR mediated genome insertion at a nuclease-induced DNA DSBs (33;34;47;48), highlighting the utility of inhibiting NHEJ in the genome engineering field. Targeting later steps in the pathway, including DNA-PK kinase activity and ligase IV activity, have been pursued with chemical inhibitors and can reduce NHEJ (34;35). Interestingly, multiple groups have also reported negligible effects of the ligase IV inhibitor, SCR7, on genome engineering experiments (49;50), highlighting the need for better, more potent and specific agents that can impact HDR mediated gene insertion. Inhibiting the initial molecular event in the NHEJ pathway, Ku interactions with DNA ends, can efficiently block NHEJ catalyzed repair and drive the processing enzymes to allow HDR mediated recombination with the appropriate donor DNA molecule. The advantage of targeting Ku is that the DSB remains unprocessed and eligible for HDR engagement and thus represents a true increase in the number of cells that are capable of HDR activity. Therefore, cells that have not been repaired by NHEJ and have not engaged the HDR system are destined for cell death, resulting in an increased ability to identify and propagate cells with the desired genetically modified sequence. Results in Figure 8 show that Ku-DBi increases HDR mediated CRISPR gene insertion of a large donor molecule by 6-fold as measured by accurate recombination results. This result shows that inhibiting Ku-DNA binding will increase targeting efficiency in genome engineering experiments at least 4-fold over other existing enhancement methods, including chemical inhibition of DNA-PKcs and Ligase IV. An additional obstacle in the genome engineering field centers around the erroneous nature of NHEJ that results in small insertions and deletions at DNA DSBs in genomic DNA. This is the driving mechanism behind off-target effects observed with engineered nucleases used for precision genome engineering (51). It is well understood that random insertion of a dsDNA molecule, a technique utilized for generating stably transfected cell lines for decades, is driven by NHEJ (48;52–55). This off-target gene insertion can also result in an increased number of non-edited cells within a larger population of cells that have been treated with CRISPRs, resulting in at times drastically larger number of samples for a researcher to screen or analyze for a cell with a specific edit. Inhibiting NHEJ should decrease these non-specific editing events. The lower number of overall puromycin resistant clonal expansions observed in the Ku-DBi treated cells (Figure 8) is likely a result of a decrease in off-target insertion of the donor molecule, activity that is typically driven by NHEJ activity. In the absence of NHEJ, we would expect to see a decrease in off-target insertion of the exogenous donor molecule. Use of chemical inhibitors targeting Ku for use in genome engineering have the additional advantage of being reversible, a benefit when optimizing CRISPR mediated gene editing experiments as the genetic background of the cells being used is unchanged. This also reduces any potential toxicity concerns that may arise within a system, as the compound is reversible and can be removed before any cell viability effects occur. In general use of these molecules for enhancing genome engineering is applicable to a wide range of cellular and model systems.

Knowledge of DNA repair pathway enzymology and the mechanism employed to respond to DNA damage has led to a series of therapeutics that have transformed clinical cancer care. The rationale for targeting DNA-PK in cancer therapy is based on its biological roles in DNA metabolism which have been elucidated from a myriad of in vitro and in vivo data, clinical analysis of patient data and our published and preliminary studies DNA-PK plays a critical role in NHEJ and the DDR pathway, combined with more recent evidence implicating it in other cellular functions including telomere progression (56;57) and cell cycle progression (58) all suggest that it plays a crucial role in cellular survival and proliferation. These biologic functions of DNA-PK/Ku provide a strong rationale for DNA-PK/Ku targeted therapy. In addition, DNA-PKcs is dysregulated in many cancers and tumors become reliant on increased DNA-PK activity to respond to the genomic instability associated with tumor cell growth (59). The enhanced DSB repair proficiency is also a major contributor to chemo- and radiotherapy resistance. Blocking DNA-PK dependent repair has been shown to increase sensitivity to various anti-cancer therapeutics and DNA-PK inhibitors are currently being assessed in combination with chemotherapy and radiotherapy and as single agents in certain DNA repair deficient cancers. This rationale is also based on the fact that DNA-PK activation and gene expression have been shown to be elevated in a number of tumor types, including ovarian cancer and hepatocellular cancer (HCC), and correlate with a poor outcome to standard therapies. Recent development of more specific and potent inhibitors has resulted in FDA approval to initiate Phase I clinical trials as a potential cancer therapeutic (60). Even with these successes moving forward in the clinic, active site targeted agents have limitations including specificity due to conserved catalytic mechanisms across kinase families, similar structural features of active sites, and the high intracellular ATP concentrations relative to the cellular concentrations of kinase inhibitors. Kinase activity, however, remains only one of the biochemical activities of the DNA-PK complex with binding DNA and supporting protein-protein interaction being integral to its role in repair and DDR. Inhibiting this activity by disrupting the Ku-DNA interaction represents a therapeutic avenue that avoids the problems associated with inhibitors targeting the active site of kinases. Results presented here definitively show increased sensitization to the DNA damaging agents bleomycin, etoposide and IR and represent a viable class of molecules for both single agent activity and enhancement of the efficacy of DNA damaging agents in the clinic.

To date, the only record of an inhibitor targeted to the Ku protein results from an in silico screen of a commercial library (61) and identifies a core scaffold that may useful in generating new molecules. However, their best compound inhibits only in the mid-micromolar range and its ability to block NHEJ catalyzed DNA DSB repair is not documented to date. In comparison, our core structure is completely different (Figure 1 and Table 1) and our synthetic medicinal chemistry approach has resulted in the synthesis of drug-like molecules with high potency and specificity towards Ku and excellent chemical properties including solubility and stability (data not shown). Studies reported here support a model where the molecules inhibit the Ku-DNA interaction by directly binding to the Ku protein and competitively blocking its association with DNA. Additionally, results in Ku deficient cell lines not only validate specificity but also indicate that cellular effects are a function of Ku inhibition. Together these results support the development of a novel class of molecules that potently inhibit NHEJ through directly disrupting the Ku-DNA interaction. The utility of such molecules for both basic science studies as well as therapeutic and commercial development is substantial.

Supplementary Data are available at NAR online.

## ACKNOWLEDGEMENT

The authors thank the IU Simon Comprehensive Cancer Center Chemical Genetics Core Facility and Pat Smith for assistance with flow cytometry.

