## Supplemental Information for "Discovery and Development of Novel DNA-PK Inhibitors by Targeting the unique Ku-DNA Interaction"

### 1) Synthetic Experimental Details:

**General.** All chemicals used for synthesis were purchased from Aldrich, Acros, Fisher Scientific and Combi-Blocks Chemical Co. (USA) and used without further purification. Anhydrous solvents were obtained from Fisher Scientific or Aldrich and used directly. All reactions involving air- or moisture-sensitive reagents were performed under a nitrogen atmosphere. $^{1}$H NMR spectra were recorded at 300 MHz using Bruker AV NMR spectrometer. $^{13}$C NMR spectra were recorded at 75 MHz using Bruker AV NMR spectrometer. The chemical shifts were reported as δ ppm relative to TMS, using the residual solvent peak as the reference unless otherwise noted. All coupling constants (*J*) are given in hertz. Data are reported as follows: chemical shift, multiplicity (s = singlet, d = doublet, t = triplet, q = quartet, p = pentet or quintet, brs = broad singlet, m = multiplet, dd = doublet of doublets, dt = doublet of triplets), number of protons and coupling constants. Thin layer chromatography was performed using Merck silica gel 60 F-254 thin layer plates, which were developed using one of the following techniques: UV fluorescence (254 nm), alkaline potassium permanganate solution (0.5% w/v) or ninhydrin (0.2% w/v) and iodine vapors. Automated flash column chromatography was carried out on prepacked silica cartridges using the indicated solvent system on Biotage Isolera chromatography system. Target compounds **68**, **149**, **322** and **245** were crystallized in 2-10% Ethanol in EtOAc, solid was collected, washed with EtOAc and then hot solutions of 20−30% EtOAc in hexanes to afford red solids. If necessary, the products were purified with automated flash column chromatography. The chemical purity of compound **68**, **149**, **322** and **245** were determined by analytical HPLC coupled to electrospray ionization mass spectrometry (LC/ESI-MS) using the area percentage method on the UV trace recorded at a wavelength of 214 nm, and compounds were found to have ≥95% purity unless otherwise specified. LC–MS analyses and purity data of compounds were obtained using an Agilent 6545 Q-ToF LC/MS instrument connected to an Agilent 1200 HPLC system, and both instruments were connected to an Agilent photodiode array (PDA) UV detector. A C-18 reversed phase column (Vydac monomeric/Phenomenex/Kinetex 2.6 μm XB-C18, 50 mm x 4.6 mm) was used as stationary phase, and water and methanol/acetonitrile (both containing 0.1−0.25% TFA) were used as mobile phase (gradient 0−100% methanol, flow 0.8 mL/min, run time 15 min). UV absorbance at the fixed wavelength of 254 nm and positive and negative ESI-MS data were recorded. The retention time and corresponding ESI-MS data were used to identify molecules. HRMS data were obtained using Waters/Macromass LCT electrospray ionization (ESI) on a time-of-flight (TOF) mass spectrometer at the Mass Spectrometry Facility at Indiana University Chemistry Department (http://msf.chem.indiana.edu).

All final compounds were purified by recrystallization or automated flash column chromatography, and the analytical and spectroscopic data confirmed their purity and structures, as detailed in the experimental procedures.

**i) General Synthetic Scheme for Synthesis of Target Compound 68, 149 and 322:**

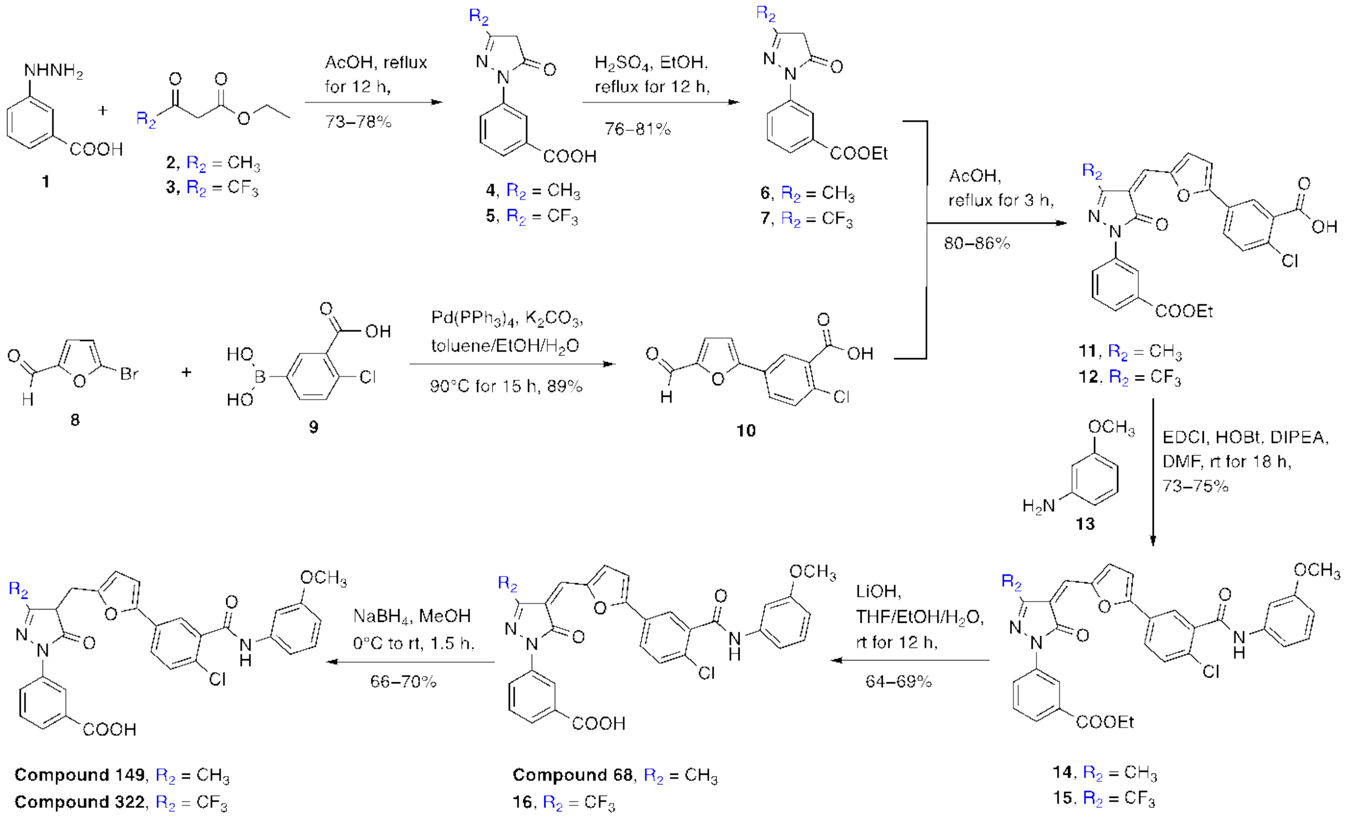

Synthesis of 3-(3-Methyl-5-oxo-4,5-dihydro-1*H*-pyrazol-1-yl)benzoic acid (**4**): Ethyl acetoacetate **2** (0.25 mL, 1.2 equiv) was added to a solution of 3-hydrazinobenzoic acid **1** (250 mg, 1 equiv) in glacial acetic acid (4 mL) under an argon atmosphere. After addition, the reaction mixture was heated at reflux with stirring for 12 h. Once the reaction was allowed to cool to room temperature, the reaction mixture was concentrated in vacuo resulting in the formation of a precipitate. The solid was filtered and washed with 5% MeOH in DCM (two times) and then two times with DCM to obtain **5** as an off-white solid (262 mg, 73% yield, requires no further purification). TLC: 4% MeOH in DCM, $R_f$ = 0.42; visualized with UV. $^1$H NMR (500 MHz, DMSO): $\delta$ 13.22 (brs, 1H, *COOH*), 8.36 (s, 1H), 8.05 (d, 1H, *J* = 8.5 Hz), 7.94 (d, 1H, *J*= 8.0 Hz), 7.70 (t, 1H, *J* = 8.0 and 16 Hz), 5.97 (s, 1H), 2.45 (s, 3H, $CH_3$). $^{13}$C NMR (125 MHz, DMSO): δ 166.48, 158.79, 154.09, 150.10, 144.90, 136.81, 132.15, 129.96, 127.57, 124.16, 120.66, 104.64, 102.19, 19.12, 14.25. HRMS (ESI): calcd for $C_{11}H_{11}N_2O_3$ [M + H]$^+$ *m*/*z* = 219.0770, found 219.0768.

Synthesis of 3-(5-Oxo-3-(trifluoromethyl)-4,5-dihydro-1*H*-pyrazol-1-yl)benzoic acid (**5**): Compound **5** was prepared by an above-described procedure using 3-hydrazinobenzoic acid **1** (250 mg, 1 equiv) and ethyl 4,4,4-trifluoroacetoacetate **3** (0.288 mL, 1.2 equiv) as starting materials. Light brown color solid (348 mg, 78% yield, requires no further purification). TLC: 4% MeOH in DCM, $R_f$ = 0.46; visualized with UV. $^1$H NMR (300 MHz, DMSO): $\delta$ 13.03 (brs, 1H, *COOH*), 8.30 (s, 1H), 8.03 (d, 1H, J = 8.13 Hz), 7.94 (d, 1H, J = 7.74 Hz), 7.66-7.61 (t, 1H, J = 7.95 and 15.87 Hz), 5.96 (s, 1H). $^{13}$C NMR (75 MHz, DMSO): $\delta$ 167.06, 154.56, 138.44, 132.23, 130.09, 128.15, 126.35, 122.72, 86.27. MS (ESI) *m*/*z* = 271.1 [M – H]$^-$.

Synthesis of Ethyl 3-(3-methyl-5-oxo-4,5-dihydro-1*H*-pyrazol-1-yl)benzoate (**6**): To a stirred suspension of 3-(3-methyl-5-oxo-4,5-dihydro-1*H*-pyrazol-1-yl)benzoic acid **4** (250 mg) in anhydrous ethanol (4 mL) was added a catalytic amount of concentrated sulfuric acid (0.3 mL) slowly under an argon atmosphere. The reaction mixture was refluxed for 12 h, and then it was allowed to cool to room temperature. The solvent was removed under vacuum, and the obtained residue was dissolved in ethyl acetate and washed successively with saturated $NaHCO_3$ (2 × 10 mL), water, and brine solution. The organic layer was dried over $Na_2SO_4$ and concentrated under reduced pressure. The crude residue was purified by Biotage automated flash column chromatography using 0–50% EtOAc in hexanes as the eluent to furnish ethyl 3-(3-methyl-5-oxo-4,5-dihydro-1*H*-pyrazol-1-yl)benzoate **6** as a red oil (214 mg, 76% yield). TLC: 45% EtOAc in hexanes, $R_f$ = 0.44; visualized with UV. $^1$H NMR (300 MHz, $CDCl_3$): $\delta$ 8.41 (s, 1H), 8.05 (d, 1H, *J* = 8.2 Hz), 7.78 (d, 1H, *J* = 8.0 Hz), 7.38 (t, 1H, *J* = 7.95 and 15.99 Hz), 4.35–4.28 (q, 2H, $OCH_2$), 3.37 (s, 2H, $CH_2$), 2.11 (s, 3H, $CH_3$), 1.33 (t, 3H, *J*= 7.11 and 14.25 Hz, $CH_3$). $^{13}$C NMR (75 MHz, $CDCl_3$): δ 170.70, 166.14, 156.84, 138.18, 131.22, 128.82, 125.77, 122.69, 119.46, 61.10, 43.02, 16.93, 14.29. MS (ESI) *m*/*z* = 247.1 [M + H]$^+$.

Synthesis of Ethyl 3-(5-oxo-3-(trifluoromethyl)-4,5-dihydro-1H-pyrazol-1-yl)benzoate (**7**): Compound **7** was prepared by an above-described procedure using **5** (300 mg) as a starting material. Off-white solid (268 mg, 81% yield). TLC: 50% EtOAc in hexanes, $R_f$ = 0.48; visualized with UV. $^1$H

NMR (300 MHz, $CDCl_3$): *δ* 10.26 (brs, 1H, OH), 8.49 (s, 1H), 8.03-7.99 (m, 2H), 7.57-7.52 (t, 1H, *J* = 8.01 and 15.99 Hz), 5.87 (s, 1H), 4.53–4.45 (q, 2H, *$OCH_2$*), 1.47 (t, 3H, *J* = 7.14 and 14.28 Hz, *$CH_3$*). $^{13}C$ NMR (75 MHz, $CDCl_3$): *δ* 167.90, 152.81, 138.30, 130.37, 129.52, 128.23, 127.38, 122.76, 86.75, 62.41, 14.29. MS (ESI) *m*/*z* = 301.1 $[M + H]^+$.

Synthesis of 2-Chloro-5-(5-formylfuran-2-yl)benzoic acid (**10**): A solution of $K_2CO_3$ (1.18 g, 3 equiv) in water (5 mL) was added to a mixture of 5-bromo-2-furaldehyde **8** (500 mg, 1 equiv) and 4-chloro-3-carboxyphenylboronic acid **9** (687 mg, 1.2 equiv) in toluene/ethanol (1:1, v/v, 25 mL). The mixture was degassed with argon for 5 min, and then $Pd(PPh_3)_4$ (165 mg, 0.05 equiv) was added. The reaction mixture was stirred at 90 °C for 15 h. The reaction mixture was cooled to room temperature, filtered through Celite, and washed with water (2 × 10 mL). The pH of the solution was adjusted to 1–2 by addition of 6 N HCl solution. The precipitated reaction mixture was extracted with dichloromethane (3 × 50 mL); the combined organic fractions were washed with brine, dried over anhydrous $Na_2SO_4$, and concentrated under reduced pressure. The crude product was triturated with 20–30% EtOAc in hexanes (2 times), and solid was filtered to afford 2-chloro-5-(5-formylfuran-2-yl)benzoic acid **10** (637 mg, 89% yield) as an off-white solid. TLC: 60% EtOAc in hexanes, $R_f$ = 0.40; visualized with UV and $KMnO_4$ solution. $^1H$ NMR (300 MHz, DMSO): *δ* 13.74 (brs, 1H, *COOH*), 9.63 (s, 1H, *CHO*), 8.23 (d, 1H, *J* = 2.22 Hz), 8.01 (dd, 1H, *J* = 2.28 and 8.43 Hz), 7.70 (d, 1H, *J* = 8.34 Hz), 7.67 (d, 1H, *J* = 2.85 Hz), 7.45 (d, 1H, *J* = 3.75 Hz). $^{13}C$ NMR (75 MHz, DMSO): δ 178.64, 166.63, 156.44, 152.47, 132.93, 132.78, 132.13, 129.00, 128.10, 127.23, 110.56. MS (ESI) *m*/*z* = 249.0 $[M – H]^–$.

Synthesis of (*Z*)-2-Chloro-5-(5-((1-(3-(ethoxycarbonyl)phenyl)-3-methyl-5-oxo-1*H*-pyrazol-4(5*H*) ylidene)methyl)furan-2-yl)benzoic Acid (**11**): Ethyl 3-(3-methyl-5-oxo-4,5-dihydro-1*H*-pyrazol-1-yl)benzoate **6** (200 mg, 1 equiv) and 2-chloro-5-(5-formylfuran-2-yl)benzoic acid **10** (203 g, 1 equiv) were dissolved in glacial acetic acid (10 mL). The reaction mixture was heated at reflux with stirring for 3 h. Solvent was removed in vacuo, and solid was suspended in EtOH, filtered, washed with EtOH, EtOAc, and DCM (2 times each) to obtain **11** as a red solid (311 mg, 80% yield, requires no further purification). TLC: 5% MeOH in DCM, $R_f$ = 0.45; visualized with UV. Major *Z*-isomer data: $^1H$ NMR (300 MHz, DMSO) δ 13.75 (brs, 1H, *COOH*), 8.63 (d, 1H, *J* = 3.87 Hz), 8.48 (t, 1H, *J* = 1.86 and 3.69 Hz), 8.27 (d, 1H, *J* = 2.22 Hz), 8.19 (d, 1H, *J* = 7.08 Hz), 8.01 (dd, 1H, *J* = 2.22 and 8.43 Hz), 7.76–7.64 (m, 3H), 7.56–7.51 (m, 2H), 4.36–4.29 (q, 2H, *$OCH_2$*), 2.64 (s, 0.29H, minor isomer, *$CH_3$*), 2.32 (s, 2.71H, major isomer, *$CH_3$*), 1.33 (t, 3H, *J* = 7.08 and 14.16 Hz, *$CH_3$*). $^{13}C$ NMR (75 MHz, DMSO): δ 166.59, 165.88, 162.11, 157.65, 151.64, 150.82, 138.96, 133.15, 132.67, 131.11, 130.91, 130.08, 129.77, 128.96, 127.84, 127.26, 125.17, 122.41, 121.60, 118.43, 112.91, 61.38, 14.65, 13.29. MS (ESI) *m*/*z* = 477.1 $[M – H]^–$.

Synthesis of (*Z*)-2-Chloro-5-(5-((1-(3-(ethoxycarbonyl)phenyl)-5-oxo-3-(trifluoromethyl)-1,5-dihydro-4*H*-pyrazol-4-ylidene)methyl)furan-2-yl)benzoic acid (**12**): Compound **12** was prepared by an above-described procedure using **7** (250 mg, 1 equiv) and **10** (208 mg, 1 equiv) as starting materials. Red solid (382 mg, 86% yield). TLC: 5% MeOH in DCM, $R_f$ = 0.48; visualized with UV. Major *Z*-isomer data: $^1$H NMR (300 MHz, DMSO) δ 13.73 (brs, 1H, *COOH*), 8.73 (s, 1H), 8.40 (d, 2H, *J* = 19.05 Hz), 8.11 (t, 2H, *J* = 9 and 17.7 Hz), 7.82 (d, 1H, *J* = 7.77 Hz), 7.74 (s, 1H), 7.69-7.57 (m, 3H), 4.36–4.29 (q, 2H, $OCH_2$), 1.36 (t, 3H, *J* = 7.08 and 14.19 Hz, $CH_3$). $^{13}$C NMR (75 MHz, DMSO): δ 166.56, 165.58, 161.24, 160.38, 150.71, 138.10, 133.99, 132.98, 131.73, 131.08, 130.02, 129.46, 127.88, 127.29, 126.76, 123.83, 119.75, 114.40, 114.08, 61.52, 14.60. MS (ESI) *m*/*z* = 531.1 $[M – H]^-$.

Synthesis of (*Z*)-Ethyl 3-(4-((5-(4-Chloro-3-((3-methoxyphenyl)carbamoyl)phenyl)furan-2-yl)methylene)-3-methyl-5-oxo-4,5-dihydro-1*H*-pyrazol-1-yl)benzoate (**14**): To a solution of compound **11** (300 mg, 1 equiv) in dry DMF (6 mL) were added EDCI·HCl (180 mg, 1.5 equiv), HOBt (127 mg 1.5 equiv), and DIPEA (0.16 mL, 1.5 equiv), and the mixture was stirred for 30 min at room temperature under an argon atmosphere. *m*-Anisidine **13** (73 µL, 1.05 equiv) and DIPEA (0.16 mL, 1.5 equiv) were added to the reaction mixture. The reaction mixture was stirred at room temperature for 18 h. The reaction mixture was poured into water and extracted with EtOAc (3 × 20 mL). The combined organic extracts were washed with saturated $NaHCO_3$ (2 × 10 mL), brine, dried over $Na_2SO_4$, and concentrated under reduced pressure. The product was triturated with mixture of EtOAc in hexanes (2–3 times) to afford **14** (267 mg, 73% yield) as a red solid. TLC: 3% MeOH in EtOAc, $R_f$ = 0.47; visualized with UV. Major *Z*-isomer data: $^1$H NMR (300 MHz, DMSO) δ 10.64 (s, 1H, *NH*), 8.65 (d, 1H, *J* = 3.81 Hz), 8.52 (t, 1H, *J* = 1.83 and 3.63 Hz), 8.22–8.14 (m, 2H), 8.06 (dd, 1H, *J* = 2.16 and 8.46 Hz), 7.81–7.70 (m, 3H), 7.64–7.53 (m, 2H), 7.43 (s, 1H), 7.29–7.27 (m, 2H), 6.74–6.70 (m, 1H), 4.37–4.30 (q, 2H, $OCH_2$), 3.75 (s, 3H, $OCH_3$), 2.70 (s, 0.58H; minor isomer, $CH_3$), 2.33 (s, 2.42H; major isomer, $CH_3$), 1.33 (t, 3H, *J* = 7.11 and 14.19 Hz, $CH_3$). $^{13}$C NMR (75 MHz, DMSO): δ 165.91, 164.69, 162.20, 160.01, 157.96, 151.75, 150.82, 140.40, 139.00, 138.22, 131.78, 131.52, 131.21, 130.66, 127.96, 125.69, 125.28, 121.63, 118.58, 112.32, 109.89, 105.83, 61.42, 55.51, 14.66, 13.30. MS (ESI) *m*/*z* = 584.1 $[M + H]^+$. HRMS (ESI): calcd for $C_{32}H_{26}N_3O_6Cl$ $[M]^+$*m*/*z* = 583.1510, found 583.1520.

Synthesis of Ethyl (*Z*)-3-(4-((5-(4-chloro-3-((3-methoxyphenyl)carbamoyl)phenyl)furan-2-yl)methylene)-5-oxo-3-(trifluoromethyl)-4,5-dihydro-1*H*-pyrazol-1-yl)benzoate (**15**): Compounds **15** was prepared by an above synthetic procedure described for the preparation of amide **14** using compound **12** (300 mg) as a starting material. Red solid (269 mg, 75% yield). TLC: 3% MeOH in DCM, $R_f$ = 0.52; visualized with UV. Major *Z*-isomer data: $^1$H NMR (300 MHz, DMSO) δ 10.53 (brs, 1H, *NH*), 8.48 (t, 1H, *J* = 1.83 and 3.66 Hz), 8.22–8.19 (d, 2H, *J* = 7.55 Hz), 7.84–7.80 (d, 2H, *J* = 7.77 Hz), 7.75 (d, 1H, *J* = 1.95 Hz), 7.66–7.52 (m, 3H), 7.40 (brs, 1H), 7.28–7.23 (m, 2H), 7.03 (d, 1H, *J* = 3.33 Hz), 6.71–6.67 (m, 1H), 4.38–4.30 (q, 2H, $OCH_2$), 3.74 (s, 3H, $OCH_3$), 1.34 (t, 3H, *J* = 7.11 and 14.19 Hz, $CH_3$). $^{13}$C NMR (75 MHz, DMSO): δ 165.83, 165.07, 159.97, 158.76, 157.58, 155.94, 150.10, 140.48, 139.86, 137.78, 130.94, 130.78, 129.80, 126.56, 125.51, 124.37, 121.30, 112.32, 109.83, 105.81, 98.53, 61.46, 55.46, 14.64. MS (ESI) *m*/*z* = 638.1 $[M + H]^+$. HRMS (ESI): calcd for $C_{32}H_{23}F_3N_3O_6ClNa$ $[M + Na]^+$*m*/*z* = 660.1125, found 660.1128.

Synthesis of (*Z*)-3-(4-((5-(4-Chloro-3-((3-methoxyphenyl)carbamoyl)phenyl)furan-2-yl)methylene)-3-methyl-5-oxo-4,5-dihydro-1*H*-pyrazol-1-yl)benzoic acid (**68**): To a stirred suspension of ester **14** (250 mg, 1 equiv) in THF/EtOH/$H_2O$ (4:2:1, 7 mL) was added LiOH (117 mg, 10 equiv). The reaction mixture was stirred at room temperature for 12 h. Solvent was removed in vacuo, and residue was acidified to pH 2–3 using 20% citric acid solution. The product was extracted with EtOAc (3 × 20 mL). The combined organic extracts were washed with brine, dried over $Na_2SO_4$, and concentrated under reduced pressure. The product was crystallized in 5% EtOH in EtOAc, and solid was collected, washed with EtOAc and then hot solutions of 20–30% EtOAc in hexanes to afford target compound **68** (164 mg, 69% yield) as a red solid. Major *Z*-isomer data: $^1$H NMR (300 MHz, DMSO) $\delta$ 13.04 (brs, 1H, *COOH*), 10.65 (s, 1H, *NH*), 8.69 (d, 1H, *J* = 3.16 Hz), 8.55 (t, 1H, *J* = 1.95 and 3.5 Hz), 8.31–8.19 (m, 2H), 8.08–7.97 (m, 1H), 7.80–7.70 (m, 3H), 7.65–7.55 (m, 2H), 7.43 (s, 1H), 7.29–7.28 (m, 2H), 6.74–6.69 (m, 1H), 3.76 (s, 3H, $OCH_3$), 2.73 (s, 0.51H; minor isomer, $CH_3$), 2.34 (s, 2.49H; major isomer, $CH_3$). $^{13}$C NMR (75 MHz, DMSO): $\delta$ 172.50, 167.51, 167.28, 165.08, 164.69, 162.22, 160.01, 157.94, 151.59, 140.41, 138.93, 132.03, 131.93, 131.22, 130.72, 129.72, 129.47, 125.48, 124.69, 122.30, 121.74, 112.32, 109.89, 105.83, 55.52, 13.30. MS (ESI) *m*/*z* = 554.1 [M – H]$^-$. HRMS (ESI): calcd for $C_{30}H_{21}N_3O_6Cl$ [M – H]$^-$ *m*/*z* = 554.1119, found 554.1122. HPLC purity: 97.72%.

<u>Synthesis of (*Z*)-3-(4-((5-(4-Chloro-3-((3-methoxyphenyl)carbamoyl)phenyl)furan-2-yl)methylene)-5-oxo-3-(trifluoromethyl)-4,5-dihydro-1*H*-pyrazol-1-yl)benzoic acid (**16**)</u>: Compound **16** was prepared by an above hydrolysis procedure described for the preparation of compound **68** using compound **15** (250 mg) as a starting material. Product was purified using 2–5% MeOH in DCM (1% AcOH in DCM) solvent system on automated flash column chromatography. Red solid (153 mg, 64% yield). Major *Z*-isomer data: $^1$H NMR (300 MHz, DMSO) $\delta$ 10.56 (brs, 1H, *NH*), 8.49 (s, 1H), 8.20–8.17 (d, 2H, *J* = 7.5 Hz), 7.85–7.79 (d, 2H, *J* = 7.62 Hz), 7.75 (d, 1H, *J* = 1.68 Hz), 7.66–7.51 (m, 3H), 7.41 (brs, 1H), 7.28–7.23 (m, 2H), 7.03 (d, 1H, *J* = 3.36 Hz), 6.71–6.68 (m, 1H), 3.74 (s, 3H, $OCH_3$). $^{13}$C NMR (75 MHz, DMSO): $\delta$ 175.00, 171.76, 167.39, 165.08, 159.97, 158.74, 156.00, 149.48, 140.49, 139.80, 137.78, 131.82, 130.06, 129.97, 127.99, 126.69, 125.26, 124.40, 123.51, 121.57, 112.32, 109.84, 108.65, 105.75, 98.53, 55.46. MS (ESI) *m*/*z* = 608.1 [M – H]$^-$. HRMS (ESI): calcd for $C_{30}H_{18}F_3N_3O_6Cl$ [M – H]$^-$ *m*/*z* = 608.0836, found 608.0833.

<u>Synthesis of 3-(4-((5-(4-Chloro-3-((3-methoxyphenyl)carbamoyl)phenyl)furan-2-yl)methyl)-3-methyl-5-oxo-4,5-dihydro-1*H*-pyrazol-1-yl)benzoic acid (**149**)</u>: To a suspension of **68** (100 mg, 1 equiv.) in anhydrous methanol (5 mL) at 0°C was added sodium borohydride (20 mg, 3 equiv.) in portions. During addition gas evolution was observed, and the color of the solution changed from dark red to yellowish orange. The resulting solution was stirred at room temperature for 1.5 h. Solvent was removed *in vacuo* and residue was acidified to pH 2-3 using 20% citric acid solution. The product was extracted with EtOAc (3 x 15 mL). The combined organic extracts was washed with brine, dried over $Na_2SO_4$ and concentrated under reduced pressure. The product was crystallized in 2% EtOH in EtOAc, solid was collected, washed with EtOAc and then hot solutions of 20-30% EtOAc in hexanes to afford target compound **149** (66 mg, 66% yield) as a red solid. $^1$H NMR (300 MHz, DMSO): $\delta$ 13.12 (brs, 1H, *COOH*), 10.67 (s, 1H, *NH*), 8.69- 8.55 (m, 1H), 8.32 (s, 1H), 8.23-8.19 (m, 1H), 8.06-8.00 (m, 1H),

7.84-7.74 (m, 3H), 7.65-7.55 (m, 2H), 7.43 (brs, 1H), 7.29-7.27 (t, 2H), 6.75-6.71 (m, 1H), 3.76 (s, 3H), 2.12 (s, 3H). MS (ESI) *m*/*z* = 610.1 $[M - H]^-$. HRMS (ESI): calcd for $C_{30}H_{21}F_3N_3O_6ClNa$ $[M + Na]^+$ *m*/*z* = 634.0969, found 634.09670. HPLC purity: 96.18%.

Synthesis of 3-(4-((5-(4-chloro-3-((3-methoxyphenyl)carbamoyl)phenyl)furan-2-yl)methyl)-5-oxo-3-(trifluoromethyl)-4,5-dihydro-1*H*-pyrazol-1-yl)benzoic acid (**322**): Compound **322** was prepared by an above reduction procedure described for the preparation of compound **149** using compound **16** (90 mg) as a starting material. The product was crystallized in 2% EtOH in EtOAc, solid was collected, washed with EtOAc and then hot solutions of 20-30% EtOAc in hexanes to afford target compound **322** (63 mg, 70% yield) as a red solid. $^{1}$H NMR (300 MHz, DMSO) $\delta$ 10.54 (brs, 1H, *NH*), 8.49 (s, 1H), 8.20–8.17 (d, 2H, *J* = 7.5 Hz), 7.84–7.78 (d, 2H, *J* = 7.56 Hz), 7.75 (d, 1H, *J* = 1.84 Hz), 7.66–7.51 (m, 3H), 7.41 (brs, 1H), 7.28–7.23 (m, 2H), 6.71–6.68 (m, 1H), 3.74 (s, 3H, *OCH*$_3$). $^{13}$C NMR (75 MHz, DMSO): $\delta$ 175.00, 171.76, 167.39, 165.08, 159.96, 158.74, 156.00, 149.98, 140.49, 139.79, 137.78, 131.82, 130.05, 129.97, 128.36, 126.68, 125.09, 124.40, 124.40, 123.51, 121.57, 112.32, 109.83, 108.78, 105.75, 98.53, 55.46. MS (ESI) *m*/*z* = 556.1 $[M - H]^-$. HRMS (ESI): calcd for $C_{30}H_{23}N_3O_6Cl$ $[M - H]^-$ *m*/*z* = 556.1275, found 556.1279. HPLC purity: 98.34%.

**ii) Synthetic Scheme for Synthesis of Compound 245:**

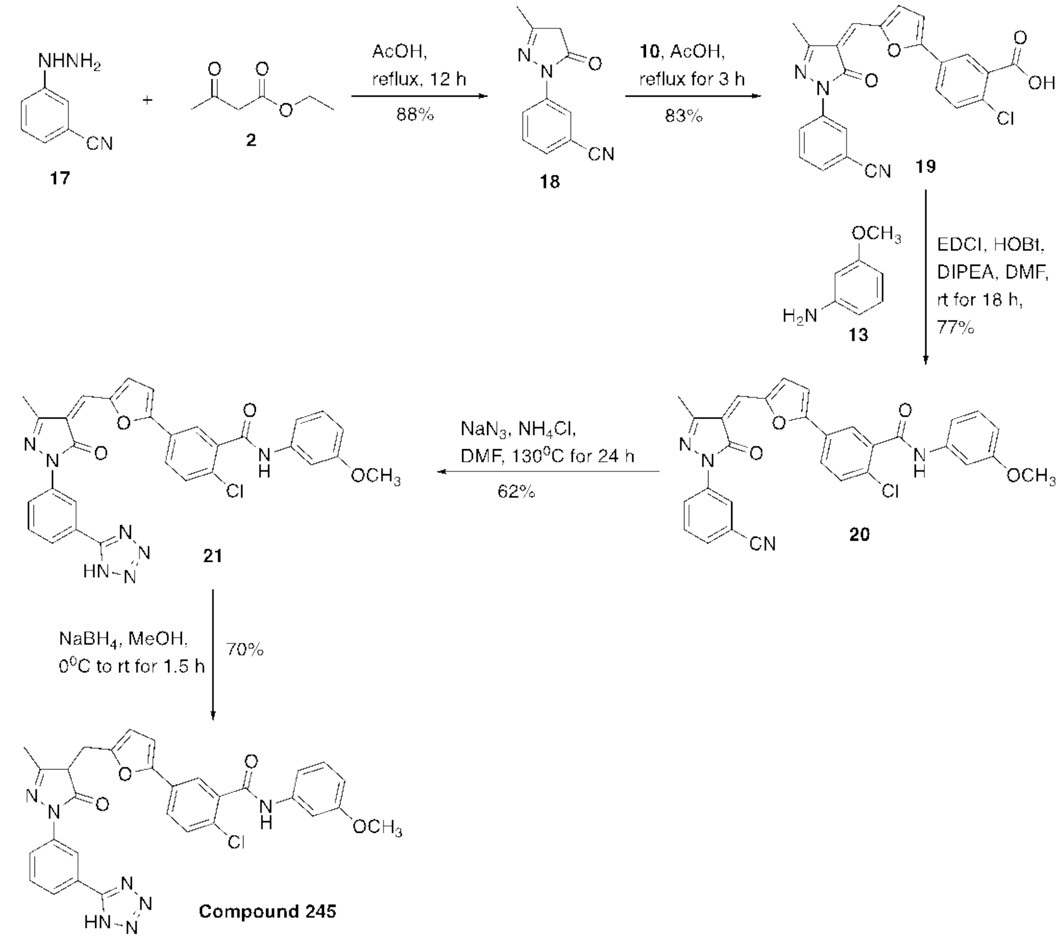

Synthesis of 3-(3-Methyl-5-oxo-4,5-dihydro-1*H*-pyrazol-1-yl)benzonitrile (**18**): Compound **18** was synthesized by an above synthetic procedure described for the preparation of intermediate **4** using 3-hydrazinobenzonitrile **17** (200 mg, 1 equiv) and ethyl acetoacetate **2** (0.228 mL, 1.2 equiv) as starting materials. Off-white solid (363 mg, 88% yield, requires no further purification). TLC: 40% EtOAc in hexanes, $R_f$ = 0.51; visualized with UV. $^1$H NMR (300 MHz, DMSO): *δ* 8.20 (s, 1H), 8.01 (d, 1H, *J* = 8.4 Hz)), 7.99-7.97 (m, 1H), 7.65-7.62 (m, 1H), 5.37 (s, 1H), 2.13 (s, 3H, $CH_3$). MS (ESI) *m*/*z* = 200.1 [M + H]$^+$.

Synthesis of (*Z*)-2-Chloro-5-(5-((1-(3-cyanophenyl)-3-methyl-5-oxo-1,5-dihydro-4*H*-pyrazol-4-ylidene)methyl)furan-2-yl)benzoic acid (**19**): Compound **19** was synthesized by an above synthetic procedure described for the preparation of intermediate **11** using compound **18** (170 mg, 1 equiv) and 2-chloro-5-(5-formylfuran-2-yl)benzoic acid **10** (214 mg, 1 equiv) as starting materials. Red solid (305 mg, 83% yield). Major *Z*-isomer data: $^1$H NMR (300 MHz, DMSO): *δ* 8.58 (d, 1H, *J* = 3.84 Hz), 8.23 (s, 1H), 8.18-8.14 (m, 2H), 7.98-7.95 (dd, 1H, *J* = 2.22 and 8.4 Hz), 7.75 (s, 1H), 7.59-7.54 (m, 4H), 2.61 (s, 0.45H; minor isomer, $CH_3$), 2.30 (s, 2.55H; major isomer, $CH_3$). $^{13}$C NMR (75 MHz, DMSO): *δ* 166.41, 162.18, 157.90, 152.06, 150.75, 139.14, 133.44, 131.25, 130.79, 128.89, 127.94, 127.57, 127.08, 122.24, 120.56, 118.98, 112.14, 13.31. MS (ESI) *m*/*z* = 430.1 [M **–** H]$^-$.

Synthesis of (*Z*)-2-Chloro-5-(5-((1-(3-cyanophenyl)-3-methyl-5-oxo-1,5-dihydro-4*H*-pyrazol-4-ylidene)methyl)furan-2-yl)-*N*-(3-methoxyphenyl)benzamide (**20**): Compounds **20** was prepared by an above synthetic procedure described for the preparation of amide **14** using compound **19** (180 mg) as a starting material. Red solid (172 mg, 77% yield). TLC: 4% MeOH in DCM, $R_f$ = 0.49; visualized with UV. Major *Z*-isomer data: Major *Z*-isomer data: $^1$H NMR (300 MHz, DMSO): *δ* 10.65 (s, 1H, *NH*), 8.63 (brs, 1H), 8.31-8.15 (m, 3H), 8.07-8.04 (dd, 1H), 7.83-7.74 (m, 2H), 7.66-7.61 (m, 3H), 7.43 (s, 1H), 7.29 (brs, 2H), 6.32 (m, 1H), 3.76 (s, 3H), 2.71 (s, 0.51H; minor isomer, $CH_3$), 2.33 (s, 2.49H; major isomer, $CH_3$). $^{13}$C NMR (75 MHz, DMSO): *δ* 170.82, 164.69 162.34, 160.01, 158.19, 152.22, 150.80, 140.39, 139.23, 138.22, 131.62, 130.93, 130.16, 128.30, 127.90, 125.74, 122.51, 121.22, 120.82, 118.99, 112.32, 112.24, 109.90, 105.82, 55.51, 13.32. MS (ESI) *m*/*z* = 537.1 [M + H]$^+$. HPLC purity: 98.43%.

Synthesis of (*Z*)-5-(5-((1-(3-(1*H*-Tetrazol-5-yl)phenyl)-3-methyl-5-oxo-1,5-dihydro-4*H*-pyrazol-4-ylidene)methyl)furan-2-yl)-2-chloro-*N*-(3-methoxyphenyl)benzamide (**21**): To a solution of nitrile **20** (150 mg, 1 equiv.) in anhydrous DMF (7.5 mL) was added sodium azide (54 mg, 3 equiv.) and then $NH_4Cl$ (45 mg, 3 equiv.). The reaction mixture was heated at 130°C for 24 h. After cooling the reaction mixture, it was poured into 40-50 mL cold water and then acidified with 1N HCl to pH ~ 2. The precipitated solid was collected by filtration, washed with water 2-3 times. The crude product was crystallized in EtOH/EtOAc mixture (1:9), solid was collected, washed with EtOAc and then hot solutions of 20-30% EtOAc in hexanes to afford tetrazole **21** (100 mg, 62% yield) as a red solid. Major *Z*-isomer data: $^1$H NMR (300 MHz, DMSO): *δ* 10.66 (s, 1H, *NH*), 8.64 (d, 1H, *J* = 3.66 Hz), 8.32 (s, 1H),

8.28-8.21 (m, 2H), 8.08-8.02 (d, 1H, *J* = 8.43 Hz), 7.84 (s, 1H), 7.77-7.74 (d, 1H, *J* = 8.49 Hz), 7.65-7.62 (m, 3H), 7.43 (brs, 1H), 7.29-7.25 (m, 2H), 6.74- 6.70 (m, 1H), 3.75 (s, 3H), 2.71 (s, 0.64H; minor isomer, $CH_3$), 2.33 (s, 2.36H; major isomer, $CH_3$). $^{13}$C NMR (75 MHz, DMSO): $\delta$ 164.68, 162.35, 160.01, 158.19, 152.23, 150.80, 140.39, 139.24, 138.24, 131.62, 130.94, 130.54, 130.16, 128.16, 127.50, 125.75, 122.53, 121.23, 120.84, 118.99, 113.04, 112.32, 112.25, 109.90, 105.82, 55.51, 13.34. MS (ESI) *m*/*z* = 580.1 [M + H]$^+$. HRMS (ESI): calcd for $C_{30}H_{23}N_7O_4Cl$ [M + H]$^+$*m*/*z* = 580.1500, found 580.1496. HPLC purity: 95.77%.

Synthesis of 5-(5-((1-(3-(1*H*-Tetrazol-5-yl)phenyl)-3-methyl-5-oxo-4,5-dihydro-1*H*-pyrazol-4-yl)methyl)furan-2-yl)-2-chloro-*N*-(3-methoxyphenyl)benzamide (**245**): Compound **245** was prepared by an above reduction procedure described for the preparation of compound **149** using compound **21** (70 mg) as a starting material. The product was crystallized in 5% EtOH in EtOAc, solid was collected, washed with EtOAc and then hot solutions of 20-30% EtOAc in hexanes to afford target compound 245 (49 mg, 70% yield) as a red solid. $^1$H NMR (300 MHz, DMSO): $\delta$ 10.62 (s, 1H, *NH*), 8.32 (s, 1H), 8.21-8.17 (m, 2H), 8.08-8.02 (s, 1H), 7.80-7.77 (m, 2H), 7.69-7.65 (m, 3H), 7.53-7.44 (m, 1H), 7.29-7.26 (brs, 2H), 6.74- 6.70 (brs, 1H), 3.75 (s, 3H), 1.98 (s, 3H). MS (ESI) *m*/*z* = 582.1 [M + H]$^+$. HRMS (ESI): calcd for $C_{30}H_{25}N_7O_4Cl$ [M + H]$^+$*m*/*z* = 582.1657, found 582.1661. HPLC purity: 97.56%.

2) 2D Interactions of compound 149 and 245 with Ku70/80:

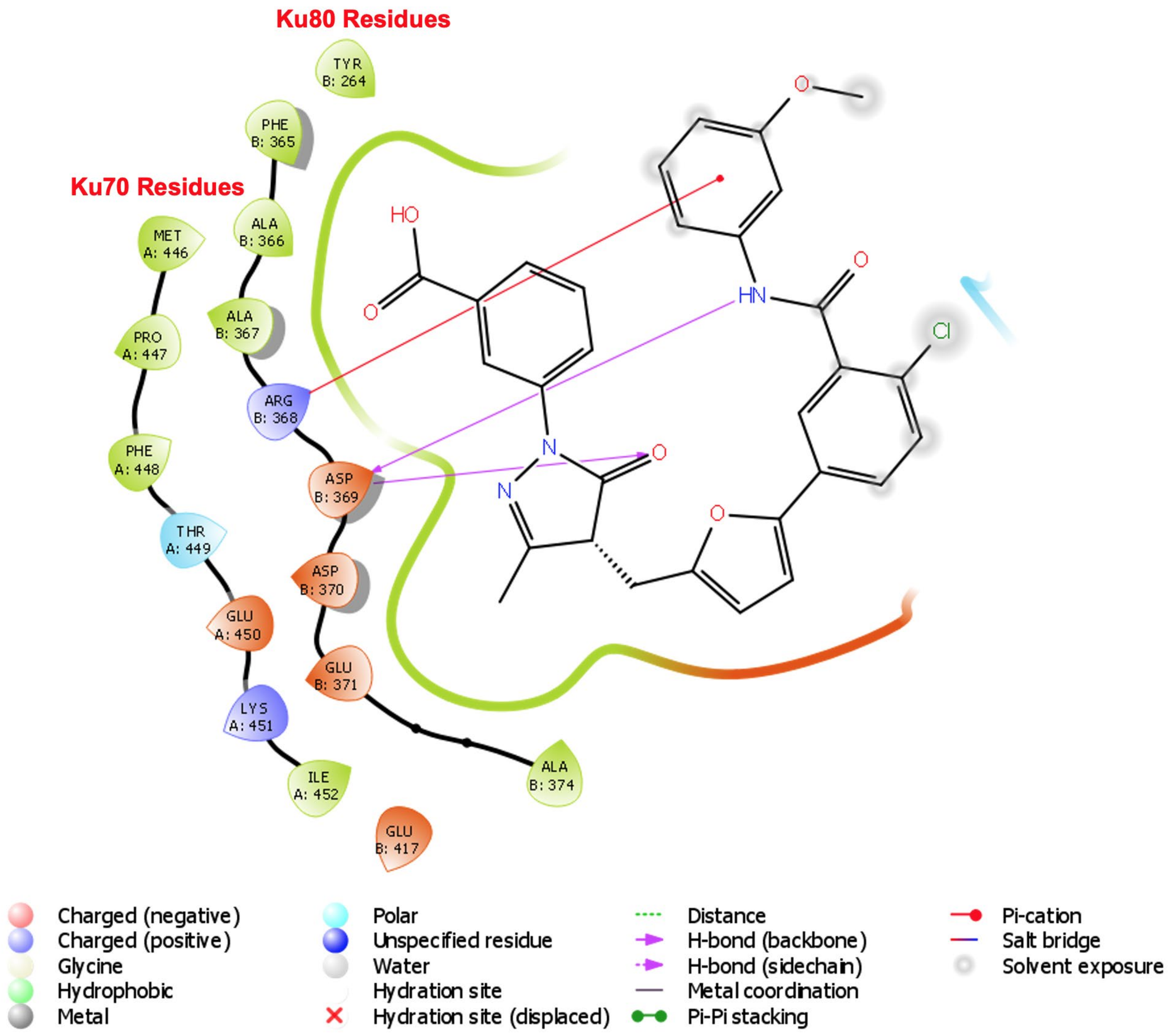

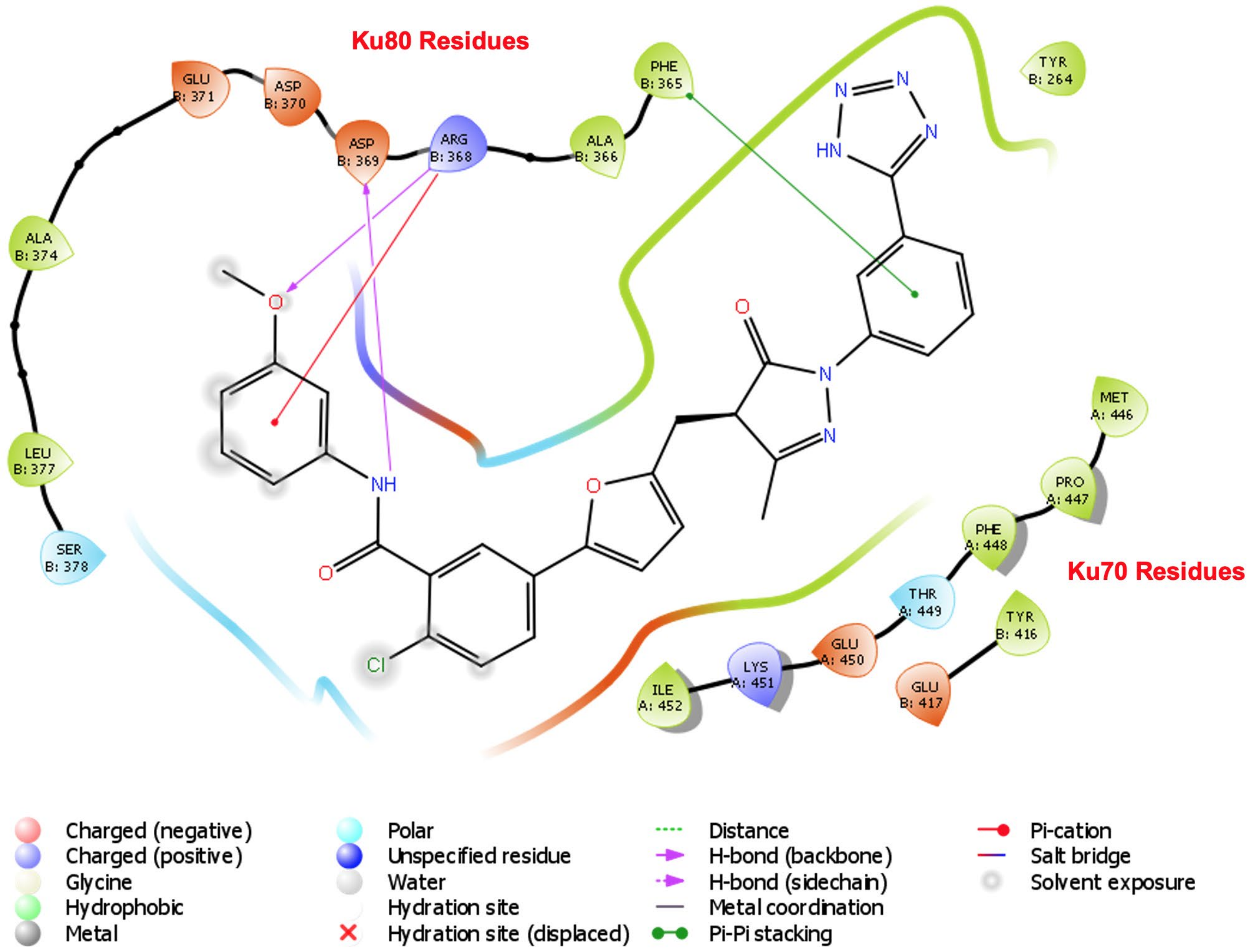
Ku80 Residues
Ku70 Residues
GLU B: 371
ASP B: 370
ASP B: 369
ARG B: 368
ALA B: 366
PHE B: 365
TYR B: 264
ALA B: 374
LEU B: 377
SER B: 378
MET A: 446
PRO A: 447
PHE A: 448
THR A: 449
GLU A: 450
LYS A: 451
ILE A: 452
TYR B: 416
GLU B: 417
Charged (negative)
Charged (positive)
Glycine
Hydrophobic
Metal
Polar
Unspecified residue
Water
Hydration site
Hydration site (displaced)
Distance
H-bond (backbone)
H-bond (sidechain)
Metal coordination
Pi-Pi stacking
Pi-cation
Salt bridge
Solvent exposure
